# Sound-mediated nucleation and growth of amyloid fibrils

**DOI:** 10.1101/2023.09.16.558053

**Authors:** Anna Kozell, Aleksei Solomonov, Roman Gaidarov, Doron Benyamin, Irit Rosenhek-Goldian, Harry Mark Greenblatt, Yaakov Levy, Ariel Amir, Uri Raviv, Ulyana Shimanovich

## Abstract

Mechanical energy, specifically in the form of ultrasound, can induce pressure variations and temperature fluctuations when applied to an aqueous media. These conditions can both positively and negatively affect protein complexes, consequently altering their stability, folding patterns, and self-assembling behavior. Despite much scientific progress, our current understanding of the effects of ultrasound on the self-assembly of amyloidogenic proteins remains limited. In the present study, we demonstrate that when the amplitude of the delivered ultrasonic energy is sufficiently low, it can induce refolding of specific motifs in protein monomers, which is sufficient for primary nucleation; this has been revealed by MD. These ultrasound-induced structural changes are initiated by pressure perturbations and are accelerated by a temperature factor. Furthermore, the prolonged action of low-amplitude ultrasound enables the elongation of amyloid protein nanofibrils directly from natively folded monomeric lysozyme protein, in a controlled manner, until it reaches a critical length. Using solution X-ray scattering, we determined that nanofibrillar assemblies, formed either under the action of sound or from natively fibrillated lysozyme, share identical structural characteristics. Thus, these results provide insights into the effects of ultrasound on fibrillar protein self-assembly and lay the foundation for the potential use of sound energy in protein chemistry.

**Significance Statement:** Understanding how and why proteins form amyloid fibrils is crucial for research into various diseases, including neurodegeneration. Ultrasound is routinely used in research settings as a tool for generating amyloid seeds (nucleation sites) from mature fibrils, which accelerate the rate of fibril growth. However, ultrasound can have various effects on aqueous media including temperature, extreme shear, and free radicals. Here we show that when the ultrasound parameters are precisely adjusted, they can be utilized as a tool for amyloid growth directly from the natively folded monomers. Thus, it is possible to induce minor changes in the folding of proteins, which trigger nucleation and accelerate amyloid growth. This knowledge lays the foundation for the potential use of sound in protein chemistry.

## Introduction

Amyloid fibrils represent a group of linear protein assemblies that possess similar characteristics, such as a common structure, fibrillation pathways, and physical properties (1). These protein assemblies are associated with dual biological functionality. On the one hand, they are linked to over 60 diseases, including neurodegenerative disorders such as Alzheimer’s and Parkinson’s (2), as well as lysozyme-related amyloidosis (3). On the other hand, structurally similar fibrillar assemblies were also found to play many functional roles in nature. Generally, amyloids are formed through misfolding or unfolding processes of proteins under specific destabilizing conditions. Factors such as pH, ionic strength, or temperature shifts can interfere with a protein’s native fold, leading to a conformational change into hydrogen (H)-bonded extended *β*-sheet-rich structures (4–7). This change coincides with a protein phase transition from a soluble monomeric state to solid nanoscale fibrils. Despite extensive research on the structure, self-assembly pathways, and biophysical properties of amyloids using various analytical techniques (8), our current understanding of the manner by which external fields affect amyloids remains limited.

External fields, particularly mechanical, like ultrasound (US), induce various physical and chemical alterations in aqueous environments (9, 10). Such alternations can result in numerous changes in fibrillar protein aggregates, including aggregate disruption, acceleration of protein aggregation, changes in protein conformation (11, 12), and enhancement of mass transfer. Since under the action of US three effects (mechanical, physical, and chemical) take place simultaneously, it is technologically challenging to separate and thus establish their contributions to the fibrillar protein aggregation process (13–16). Moreover, the effects of US on protein aggregation may be highly protein-specific, since different proteins have varying sensitivities to mechanical stress, heat, and the conformational changes induced by US. Additionally, the delivered US energy conditions, such as intensity, duration, and frequency, need to be carefully optimized to achieve the desired effect on protein species.

Recently, we comprehensively studied how sound energy affects fibrillar protein complexes, aiming to establish boundary conditions under which sound energy causes either damage (denaturation) or conversely, induces structural modifications (17). We have established that under the action of high energy, high-amplitude, and high-power ultrasound (20 kHz, 7 W), the thermal effect dominates for protein monomers and can lead to thermal protein denaturation as well as to reduced *β*-sheet content in lysozyme protein fibrils.

In the present study we observed that although a short exposure of lysozyme protein species to low-amplitude (2 W power) US did not lead to any significant structural changes, these conditions were sufficient to modify the chemical kinetics of protein fibrillation. Furthermore, long-time exposure of the monomeric protein to low-amplitude US enabled nucleation and fibril elongation in a highly controlled manner. Since long-term exposure of the aqueous solution to low-amplitude US does not lead to any significant thermal fluctuations (below the melting point of the lysozyme protein studied), it highlights the key role of pressure in modifying the self-assembling behavior of fiber-forming proteins. To gain deeper insights into the effects of the amplitude and power of US on protein monomers, we performed a comprehensive structural, morphological, and molecular dynamics (MD) analysis. Using MD, we succeeded in detecting the protein motifs, characterized by increased sensitivity to pressure and temperature fluctuations, which serve as a starting point for initiating structural changes and further fibrillation. Importantly, we observed that although the overall protein secondary structure is maintained, very small changes in the protein fold are sufficient to initiate protein-protein interaction, nucleation, and further accelerate fibril growth. Contrary to high-amplitude and high-power ultrasound, where the thermal effect dominates, with low-amplitude ultrasound, structural changes in protein species are initiated by pressure fluctuations. In summary, these results highlight the role of pressure parameters in modifying protein folding and further contribute to our understanding of the overall effects of mechanical energy in the form of ultrasound on the protein structure, conformation, and specifically on the fibrillar protein self-assembly phenomenon. The results from our study open horizons for the potential exploitation of ultrasound as a tool that can chemically modify protein complexes without changing the chemical environment.

## Results and Discussion

### Formation and growth of amyloid fibrils under low-amplitude US

Previously, we reported that short exposure of monomeric lysozyme protein to low-power US (2 W, 20% amplitude, which is equivalent to 400 J) (17), leads to the formation of small particles with a heterogeneous size distribution. The formation of the particles was accompanied by insignificant structural changes in protein, towards increasing the *β*-sheet content. These particles, however, displayed the ability to accelerate the rate of fibril growth, similarly to the seeds formed from mature fibrils via fragmentation. Interestingly, when the power of US was increased from 2 W to 7 W (equivalent to a 1000 J energy value), it led to thermal denaturation of the protein molecules. Such an observation suggests that low-power US can alter protein complexation. Since US effects in aqueous media originate from pressure fluctuations that give rise to a temperature increase (see **Figure 1 *A***), elimination or suppression of the temperature factor can open multiple routes for the non-damageable manipulation of protein-self-assembly.

**Figure 1.**
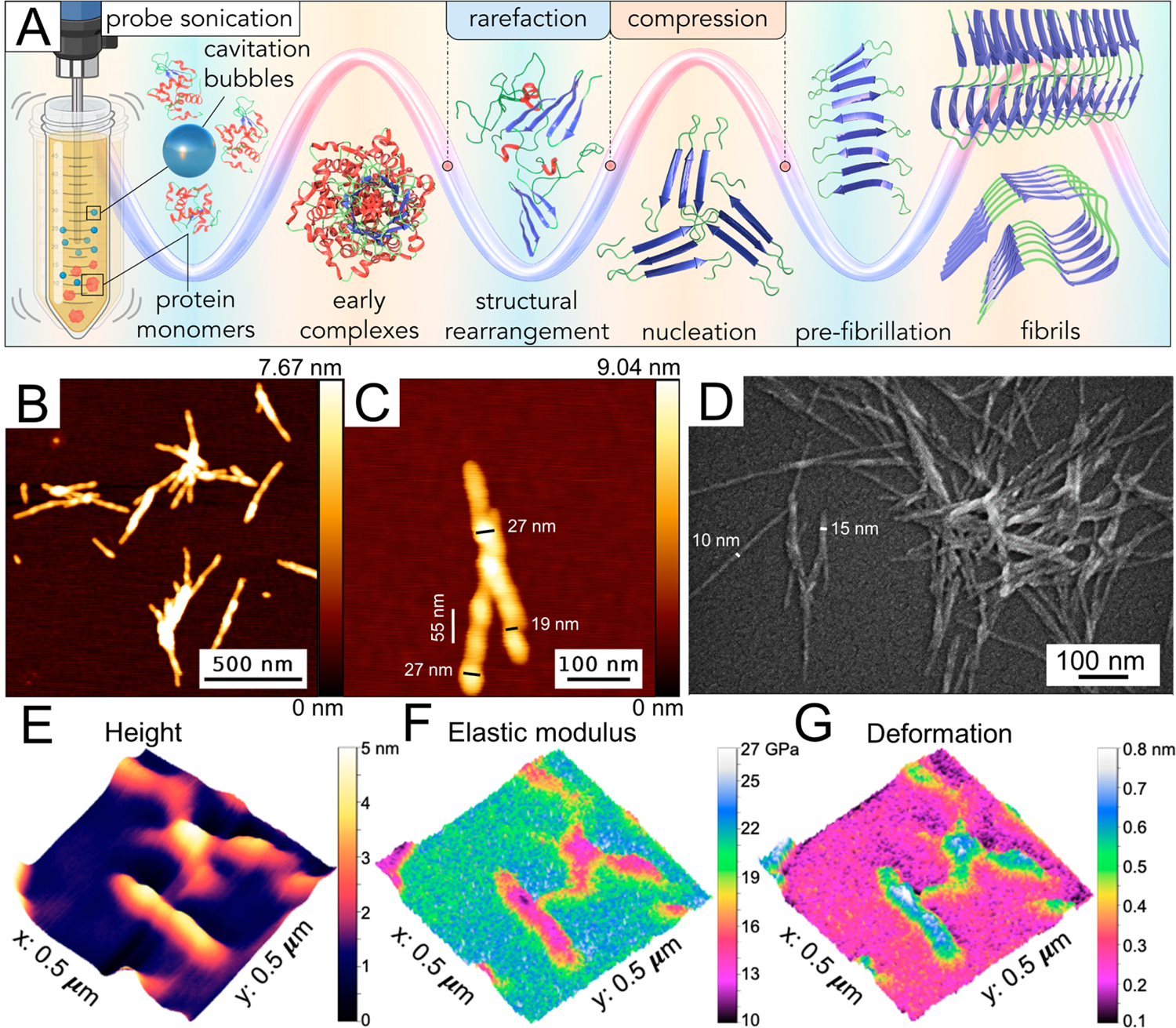
(A) Schematic representation describing amyloid fibril formation and growth under the action of low-amplitude ultrasound (see the Materials and Methods Section, “Formation of lysozyme fibrils under the action of ultrasound”). (B, C) Atomic force microscope (AFM) images of lysozyme aggregates and (D) scanning electron microscopy (SEM) images of aggregates formed by US. (E, F, G) Mechanical properties of the obtained fibrils measured by AFM nanoindentation. Separate images were prepared with biorender.com, Blender 2.93, and Autodesk Fusion 360.

To this end, we focused our current investigation on the effects of low amplitude and low power (2 W) US. First, we systematically increased the US energy acting on monomeric protein species (concentration is 1.4 mM) via maintaining a low amplitude of US wave, but we increased the exposure time from 4 min to 10 min (see the Experimental section). These conditions enabled us to eliminate thermal denaturation. Whereas for a shorter exposure time (< 8 min) we observed no changes in particle size, a relatively long (10 min) exposure of monomeric lysozyme to low-amplitude US wave leads to the formation of linear fibrillar structures of ∼10-30 nm in width, as depicted in **Figure 1 *B*** and ***C,*** and lengths varying between ∼100 nm and ∼1500 nm (**Figure 1 *D*)**. Such an observation indicates that lysozyme protein underwent nucleation and elongation after low-amplitude US exposure. The critical length for US-grown lysozyme fibrils was reached at 1000 J and with a 10 min exposure time. The growth of protein nanofibrils with US for a longer exposure time, in attempting to elongate fibrils beyond 1500 nm length, has led to fibril fragmentation, which characterizes the typical effect of US on protein fibrils, namely, mechanical fragmentation. We hypothesized that the measured critical fibril length might originate from the correlation between the mechanical characteristics of US-formed lysozyme nanofibrils and the acting shear forces. To corroborate our hypothesis, we performed a nanoindentation analysis of US-formed lysozyme nanofibrils to determine their mechanical characteristics. The nanomechanical properties are a distinguishing feature of amyloid protein fibrils. Therefore, we conducted modulus mapping on the lysozyme structures formed from monomers when exposed to ultrasound. The results from our nanomechanical analysis, summarized in **Figure 1 *F, J***, indicate that the measured rigidity of the US-formed nanofibrils is 13.75±1.50 GPa (**Figure 1 *J***). Remarkably, the value of fibril rigidity modulus is near that of fibrils formed without US exposure, which exhibited an elastic modulus of 11.95±2.51 GPa, as indicated in our earlier publication (17). Considering that ultrasound can produce pressure fluctuations varying between 0.01 and 2.5 MPa, calculated and reported in previous studies (18), such conditions do not alter protein fibril stability and enable their elongation until a critical length is reached. An interesting parameter is the nanofibril critical length, with a maximum measured value of 1500 nm. Such a value could be linked to the size of the collapsing cavity bubble formed in aqueous media by US, which varies between 0.1 and 23.7 µm in radius (18, 19).

### Morphology of amyloid fibrillar assemblies

To morphologically characterize lysozyme fibrillar assemblies formed under the action of low-amplitude US, atomic force microscopy (AFM) and scanning electron microscopy (SEM) analyses were carried out (**Figure 1 B-D**). Particles formed at 1000 J appear as short nanofibrils with dimensions of ∼8 nm in height, ∼10-30 nm in width, and lengths varying between 100 and 1500 nm. We observed the appearance of a fibril twist (twisted ribbon) when its length exceeded 200 nm. The twist repeat along the fibril axis is left-handed, with a pitch of ∼55 nm (see **Figure 1 *C*)**, which is consistent with literature reports (20–22) indicating the appearance of a similar structural organization in thermally induced aggregated lysozyme fibrils.

### Dynamics of the US-induced fibril growth

Experimentally, we find that in the presence of the ultrasound the fibrils grow to significant lengths, with a mean fibril length of ∼300 nm (Figure 2 ***B***). To better understand this phenomenon, we studied the coarse-grained, minimal models of fibril dynamics. The simplest model (referred to below as the “basic model”) includes the following three processes: the initial complexation of monomers, which further act as nucleation sites for subsequent fibril growth, elongation of pre-existing fibrils, and fragmentation of an existing fibril into two separate ones (Figure 2 ***A,*** (***1***)***-***(***3***)). In a modified model, we also considered monomer detachment from the end of the fibrils (Figure 2 ***A,*** (***4***)). We assumed first-order kinetics, implying that elongation is proportional to the free monomer concentration, and that the rate of fragmentation at any given point along the fibrils remains constant. The master equation describing the temporal dynamics is as follows:

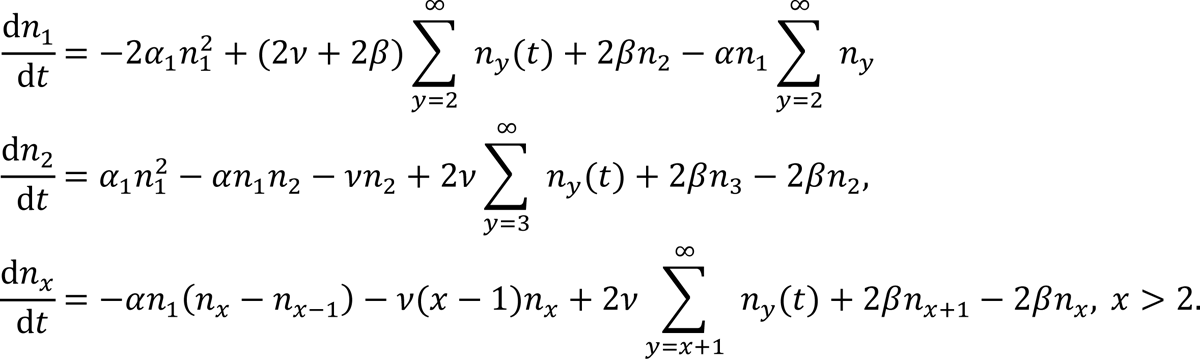

**Figure 2.**
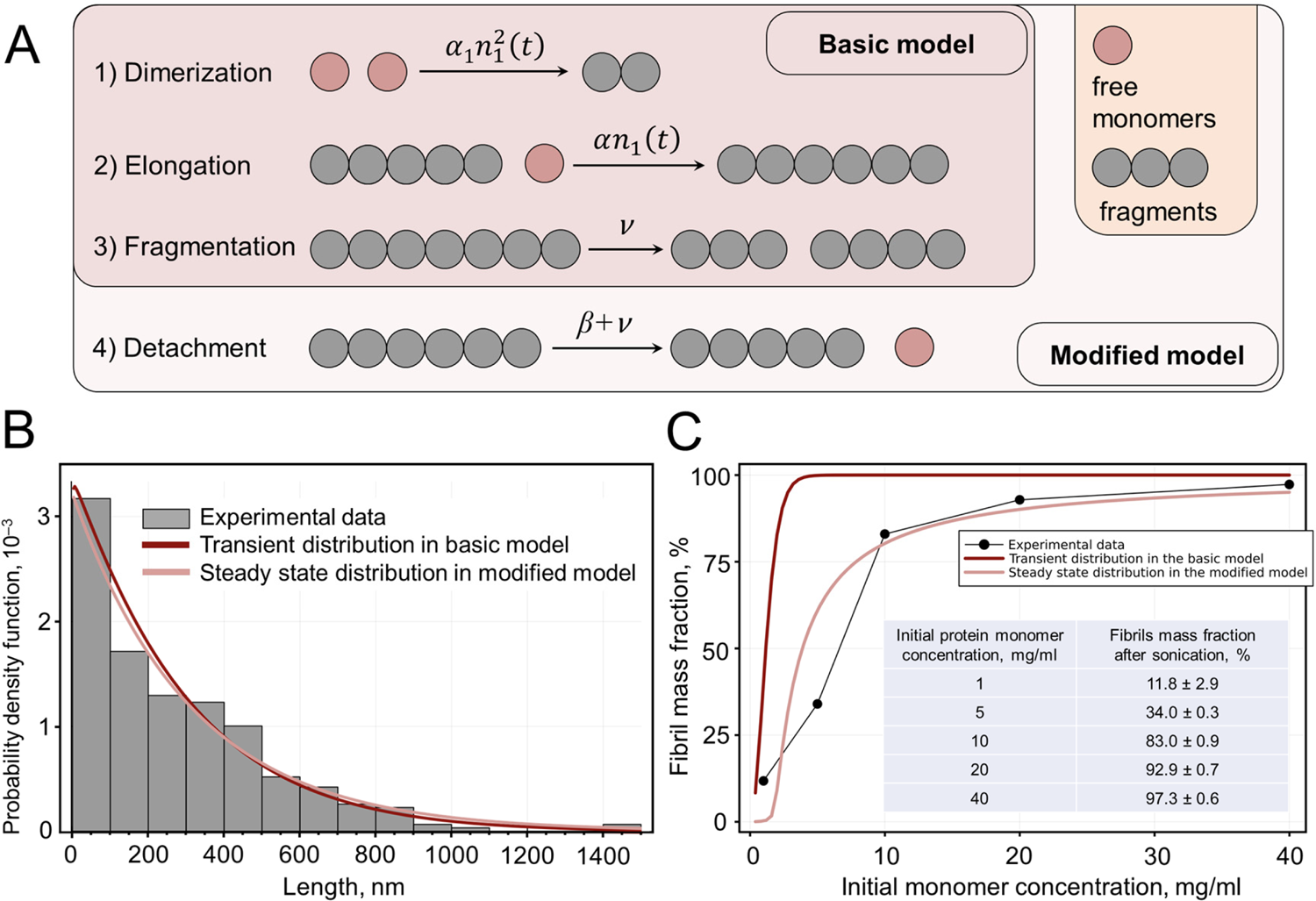
(A) A schematic representation of the model dynamics features: the pale red circles represent the free monomers and the gray structures represent fibrils (or their fragments). The arrows highlight transition processes within the system, and their respective rates are given. (B) The probability density function (PDF) of the numerical results (the red and pink lines) compared to the experimentally measured histogram (the gray bars). The basic model contains processes (1)-(3) in panel A, which can only lead to long fibrils transiently. In contrast, a modified model that contains process (4) in panel A can also lead to long fibrils in steady state. (C) Fibril mass concentration at the moment of observation (i.e., after the sonication) plotted against the initial concentration of monomers. The model curves are obtained by a numerical solution of Eq. (1) and Eq. (12), SI Appendix, with parameters reproducing the data in panel B. An inset includes a table detailing the initial protein monomer concentrations and the fraction of fibrils obtained under specific ultrasonication conditions (20 kHz, 2 W, 10 min). A photograph illustrating the samples treated by ultrasound, which vary depending on the protein concentration is presented in SI Appendix**, Fig. S2**. For a detailed explanation of the equation symbols, refer to the SI Appendix.

Here the Greek letters represent the rates of the corresponding processes (see Figure 2 ***A***), and the basic model corresponds to the case when the detachment only occurs as a part of the fragmentation process, i.e., β = 0 (see the *SI Appendix* for details).

Surprisingly, we found that within the basic model the steady-state distribution cannot explain the experimental data: in the *SI Appendix (the “Mean fibril length” section)* we proved that for any model parameters, the mean fibril length is bounded by 3, in sharp contrast to the experimental results. However, we found that the basic model can lead to a non-monotonic mean fibril length: transiently, the dynamics can lead to very long fibrils that will later become shorter as the steady-state is reached (see *SI Appendix**, Fig. S1***). The model can reproduce the experimentally measured fibril length distributions using numerical simulations for the appropriate choice of parameters (see Figure 2 ***A***).

The intuition for the failure of the basic model in producing long fibrils in steady state conditions is that in steady state the monomer pool would be depleted, and any existing long fibrils would break into smaller ones due to fragmentation; however, they would be unable to elongate since no monomers are available. We therefore hypothesized that adding an additional monomer detachment mechanism would bypass this limitation and could also provide long fibrils in steady state (process (4) in Figure 2 ***A***). Indeed, we found that the experimental data could also be explained by the modified model in steady state (Figure 2 ***B***).

Several conclusions can be drawn from these results, which suggest new avenues for future works: first, within the basic model, the observed long fibrils are a transient effect, and the fibril length distribution is out of steady state. Within the modified model, the long fibrils exist in steady state. Second, the basic model contains two key dimensionless parameters, and in order to yield long fibrils (for intermediate times), the fragmentation rate must be small compared with the elongation rate, which, in turn, must be larger than the dimerization rate. Therefore, the experimental data in combination with the physical model, yield useful constraints on the parameter space, and facilitate estimating the relevant microscopic rates in our system. Similarly, to explain the experimental data within the modified model, in addition to the previous restrictions, one must keep the detachment rate large in comparison with the fragmentation rate.

Figure 2 ***B*** shows that the experimental distribution for the fibril length is consistent with both the basic model (transiently) and the modified model (in steady state). Even though the computed distributions are similar for mature structures, there is an important distinction between them: for the modified model the computed mass fraction of free monomers, *n*_1_/*N*, is on the order of 0.1, whereas for the basic model the fraction is negligible. Thus, the experimental data on the monomer mass fraction in Figure 3 ***C*** (the dark blue bar, protein monomers) support the modified model rather than the basic model to explain the observed fibril length distribution.

**Figure 3.**
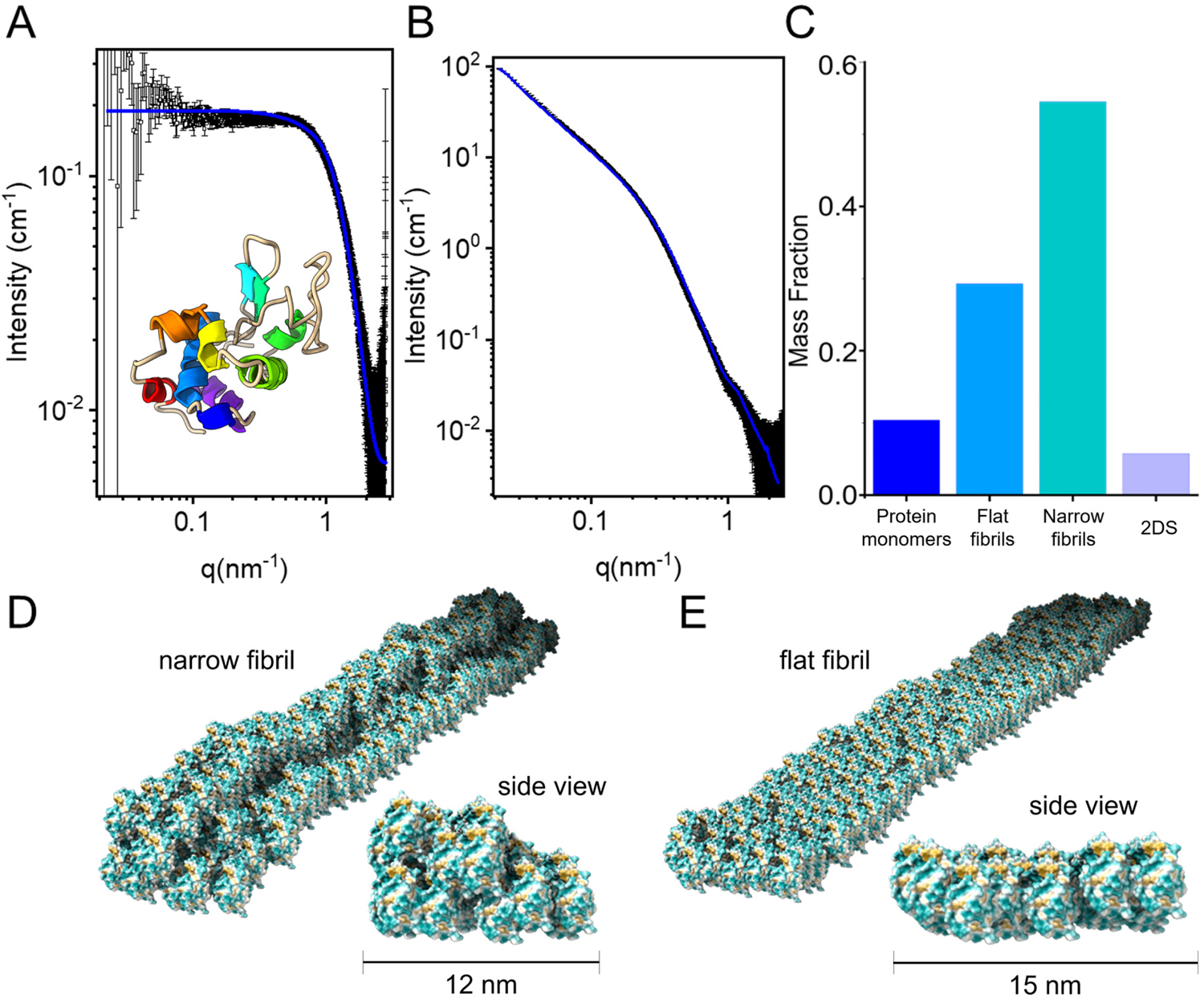
Small X-ray scattering (SAXS) analysis. (A) Azimuthally integrated background-subtracted scattering intensity *I*, as a function of *q*, the magnitude of the momentum transfer vector *q*→ (the black symbols in Figure 3 **B**) from soluble lysozyme monomers (adapted from (17) Copyright © 2023 the Author(s). Published by PNAS), (B) I vs. q from lysozyme assemblies formed after applying US irradiation to a solution of lysozyme monomers. (C) The mass fraction of the coexisting structures that formed after US irradiation (where 2DS denotes a 2D-sheet). Models of the narrow (D) and flat (E) lysozyme fibrils formed after US exposure, based on the SAXS analysis. Fibril and protein visualization: ChimeraX 1.7 (25, 26). Data analysis was performed as explained in our earlier publication (17).

To further reconfirm our findings, a set of experiments were conducted, systematically altering the initial monomer concentration parameter. Consequently, we have found that the observed dependence is better reproduced by the modified model, providing additional support to the previously discussed argument (see Figure 2 C).

### Characterization of the structural changes imposed by low-amplitude US on the protein monomers

#### Small-angle X-ray scattering (SAXS) analysis

Using solution small X-ray scattering analysis at the European Synchrotron Radiation Facility (ESRF) (see the Materials and Methods section), we investigated and compared the structures of natively folded protein (Figure 3 ***A***), natively aggregated lysozyme fibrils (17), and US-formed fibrils (Figure 3 ***B***). The data were analyzed using D+ software (23, 24). Importantly, we found that exposing natively folded lysozyme monomers to low-energy US resulted in the formation of two types of fibrillar supramolecular assemblies: flat fibrils and narrow fibrils, with mass fractions of 0.293 and 0.545, respectively Figure 3 ***C***).

The model (the blue curve in Figure 3 ***A***) was obtained by a linear combination of the monomer form-factor, M (shown in Figure 3 ***A***, modeled using PDB ID:1AKI), two fibrils (flat fibrils, denoted as FF and narrow fibrils, denoted as NF), and a 2D sheet (2DS). In the fibrils, the monomers were rotated about the z-axis by - π/9 *rad*, with respect to their orientation in PDB ID: 1AKI. To fit the data in Figure 3 ***B***, we have added the contribution of thermal fluctuations. This was done Monte Carlo simulaitons, assuming a harmonic potential between nearest-neighbor monomers with a lattice elastic constant *κ*, as previously explained (27). Using the equipartition theorem, the root mean squared displacement (RMSD) of monomers in the fibril was estimated to be 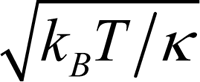 (17). In a fibril of height *H*, depth *d*, and width *W*, the location of a specific lysosome monomer, ř_ijk_ = (*x*_i_, *y*_j_, *z*_k_) was *x*_i_ = *iD*_x_, *y*_j_ = *jD*_y_, *z*_k_ = *kD*_z_, where *i*, *j*, and *k* varied between 0 and 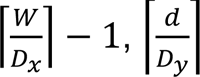 − 1, and 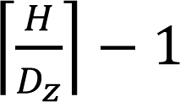, respectively. The parameters of the fibrils were *D*_x_ = *D*_z_ = 2.6 *nm, D*_y_ = 3.6 *nm, H* = 400 *nm*, *w* = 12 *nm, d* = 7 *nm*, and the value of *K* 4 ± 2 x 10^-20^ *J* ⋅ *nm*^-2^, which corresponded to an RMSD value of ∼0.3 nm, suggesting that the fibrils were rather soft, considering the Lindemann’s criterion (28). In the narrow fibril (NF), we deleted 2 (out of the 5) fibrils of the last layer of the fibril. In the flat fibril (FF), we used the same parameters with the following changes: *w* = 15.6 *nm, d* = 7.2 *nm*, and *K* = 6 ± 3 x 10^-20^ *J* ⋅ *nm*^-2^ (RMSD = 0.26 nm), and we deleted the last layer of the fibril (FF). In the 2D sheet (2DS): *H* = 400 *nm*, *w* = 300 *nm, d* = 6 *nm*, and *K* = 4 ± 2 x 10^-20^ *J* ⋅ *nm*^-2^(RMSD ∼ 0.3 nm).

Thus, our analysis revealed that low-power US triggered the formation of two types of soft nanofibrils: narrow (Figure 3 ***D***) and flat (Figure 3 ***E***), with a higher fraction of narrow fibrils. Furthermore, the narrow fibrillar shape resembles the shape of the natively self-assembled amyloid fibrils.

#### Secondary structure evaluation using Fourier transform infrared spectroscopy (FTIR) analysis of aggregated protein nanofibrils and the chemical kinetics of self-assembly

To investigate whether lysozyme nanofibrils, produced via US, display self-assembly kinetic behavior similar to amyloids, we employed the standard Thioflavin T (ThT) assay (29, 30). ThT exhibits a red shift in emission, with a measured maximum intensity at 490 nm, when it associates with amyloid fibrils enriched in *β*-sheet conformation. Recording these shifts makes it possible to accurately identify distinct stages of protein aggregation, such as the initial nucleation, fiber elongation, and the formation of secondary nucleation. We combined the US-grown nanofibrils with native lysozyme monomers in a 1:1 volume ratio. We first tested the effect of variable concentrations of untreated monomers on the kinetics of amyloid growth. As expected, the increase in monomer concentration accelerated the aggregation process, as shown in the *SI Appendix**, Fig. S3***. Furthermore, the kinetics curves (Figure 4 ***A***) indicate that the US-formed nanofibrils can accelerate kinetic growth, even at different concentrations, without any appreciable lag phase, resembling the behavior of the seeded amyloid aggregation. This supports our initial observations about the amyloid-like characteristics of the fibrils we produced by using US.

**Figure 4.**
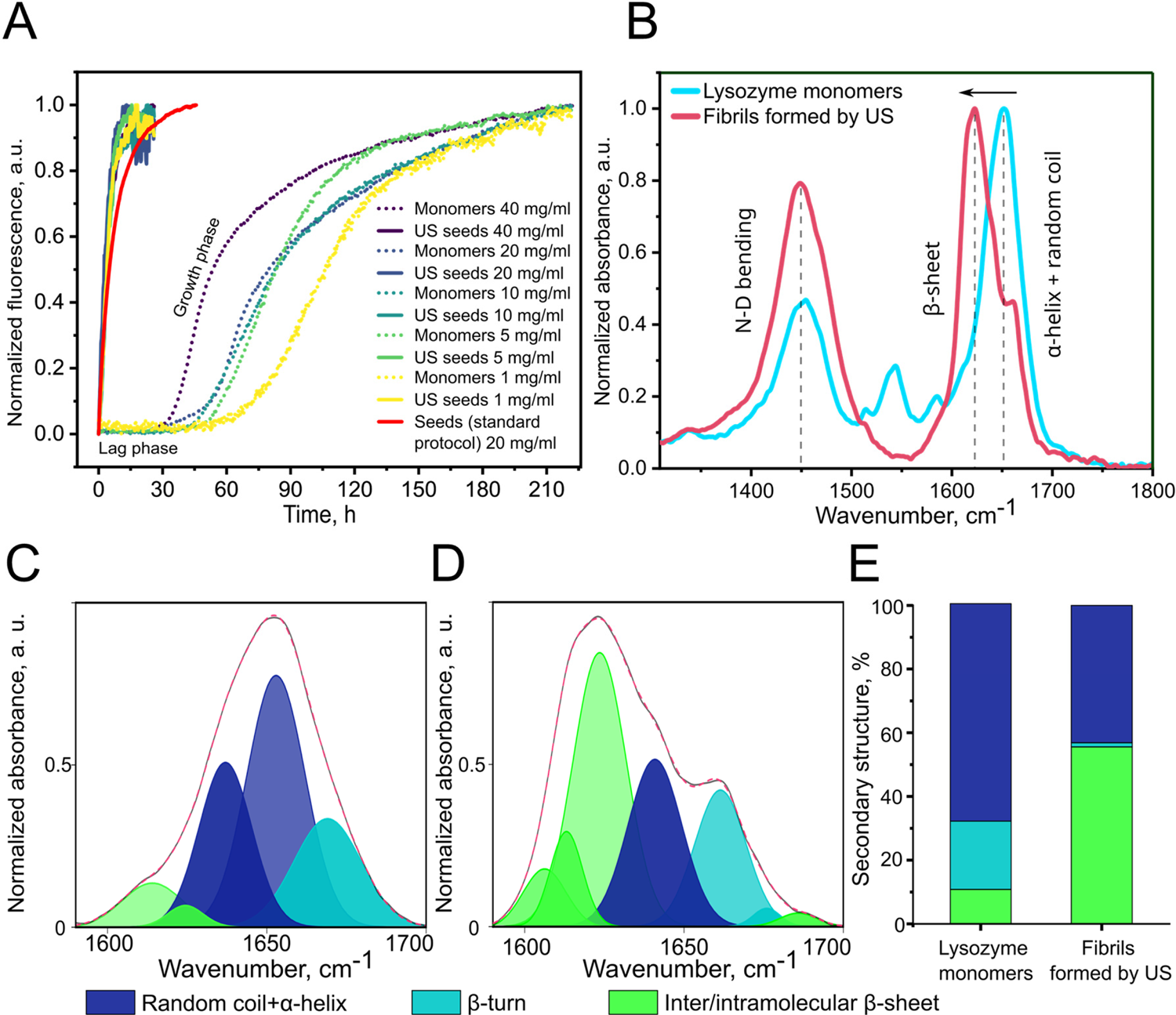
Kinetics of self-assembly and FTIR analysis of lysozyme monomers and nanofibrils formed due to low-amplitude US. (A) Analysis of the chemical kinetics of lysozyme nanofibrils formed by US at different protein concentrations (1, 5, 10, 20, 40 mg·ml^-1^) compared to seeds formed according to the standard protocol (see Experimental section), and native lysozyme monomers at the same concentrations (ratio 1:1). Non-normalized data is shown in SI Appendix, **Fig. S3**. (B) Amide I and Amide II bands of the FTIR spectra of lysozyme monomers (cyan), and a nanofibril solution formed after 10 min of US (red). (C) Deconvoluted Amide I band of lysozyme monomers, (D) nanofibril solution, (E) A comparative analysis of the secondary structure in %.

To establish the possible structural re-arrangements in fibrils that were formed by exposing lysozyme monomers to US, we conducted a secondary structure analysis by using Fourier-transform infrared spectroscopy (FTIR). In particular, we focused on the alterations in the vibration bands associated with amide I and amide II.

The experiment was performed in a deuterated solvent. Traditionally, studying the protein secondary structure using deuterated solvents, such as deuterated water (D_2_O), is a common practice in FTIR spectroscopy. Standard water can interfere with the measurements, owing to the strong absorption bands of its O-H stretching vibrations, overlapping with the proteins’ amide I and II bands. Deuterated solvents help to minimize such interference and improve the quality of the acquired spectra. D_2_O exhibits a lower absorption in the region of interest for protein secondary structure analysis, since the O-D stretching vibrations occur at different wavenumbers, compared with O-H stretching. This property enabled a clearer observation of the amide I and II bands (Figure 4 ***B***), which are crucial for evaluating the protein’s secondary structure. The aggregated amyloid structures’ vibration spectra are characterized by intermolecular *β*-sheet content (1610-1625 cm^-1^) and antiparallel amyloid *β*-sheets (1690-1705 cm^-1^) (31). These spectra, which differ from the characteristic vibrations of the native protein fold, are 1625-1635 cm^-1^ for *β*-sheet content, 1635-1665 cm^-1^ for *α*-helix and random coils, and 1665-1690 cm^-1^ for *β*-turns (Figure 4 ***C, D***).

We found that the secondary structure of the monomeric lysozyme underwent significant changes when exposed to low-amplitude ultrasound for 10 minutes. The content of β-sheet significantly increases (from 11% to 58%), adopting a conformation typical for amyloids (summarized in Figure 4 ***E***). This increase is more pronounced than the β-sheet content of 47% observed in fibrils prepared without ultrasound (17).

#### Molecular dynamics (MD) simulations

To gain further insight into the mechanism underlying lysozyme fibrillation under low-power and low-amplitude ultrasound, we performed molecular dynamics (MD) simulations of isolated lysozyme in water. We wished to model two of the forces that are present in aqueous media while applying US: temperature and pressure gradients. We focused our MD analysis on two main physical factors: (*1*) temperature, (*2*) pressure and combined when created in aqueous media upon exposure to US. We monitored changes both in the complete structure of the protein molecule and in all its secondary structure elements (*α*-helixes, *β*-sheets, and several random coils) when these forces were applied both separately and together. The model of hen egg white lysozyme (HEWL, PDB ID: 193L, Figure 5 ***A***) was used for simulations. HEWL consists of 129 amino acid residues and eight *α*-helixes (H) and two *β*-sheets (S): H1 (α-helix, 5-14, RCELAAAMKR), H2 (*α*-helix, 20-22, YRG), H3 (*α*-helix, 25-36, LGNWVCAAKFES), S1 (*β*-sheet, 43-45, TNR), S2 (*β*-sheet, 51-53, TDY), H4 (α-helix, 80-84, CSALL), H5 (*α*-helix, 89-99, TASVNCAKKIV), H6 (*α*-helix, 104-107, GMNA), H7 (*α*-helix, 109-114, VAWRNR), H8 (*α*-helix, 120-124, VQAWI).

**Figure 5.**
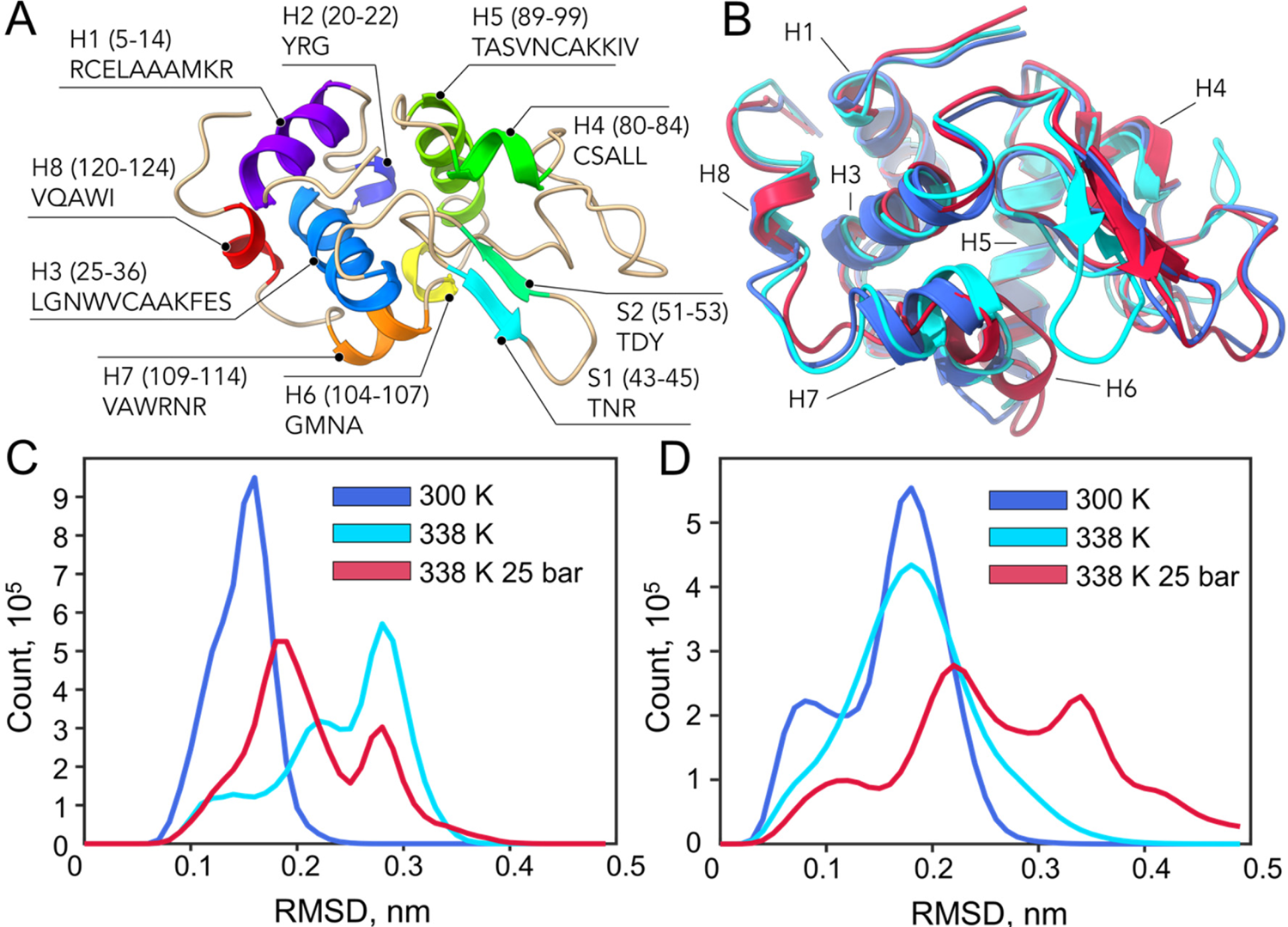
(A) Chemical structure of hen egg white lysozyme (HEWL, PDB ID: 193L, 129 amino acid residues) represented as a cartoon model. The α-helix and β-sheet structures are shown in rainbow colors and are denoted by the letters “H” and “S”, corresponding to their positions in the protein structure and the amino acid sequence. Numeration of elements started from the N-terminal; (B) the superposition of three protein structures under different conditions and different simulation time frame snapshots: blue – at 300 K, cyan – at 338 K (t = 449693 ps), and red – at 338 K 25 bar (t = 1843799 ps, state 2, see also the SI Appendix, **Fig. S19**, **Fig. S20**, **Fig. S21**, and **Fig. S22**), pointing to key structural elements; visualization: ChimeraX 1.7 (25, 26); diagrams of cumulative (0 – 3000000 ps) RMSD of C_α_ (SI Appendix, **Fig. S4**) over two runs for (C), the whole protein sequence, and (D) for H7 (see Figure 5 **A**) under different simulation conditions (referred to in the SI Appendix, **Fig. S5**, and **Fig. S14**, correspondingly).

We employed three different simulation conditions (two runs for each) for 3 microseconds recorded in 1 picosecond increments (3·10^6^ points) for three states of the protein in aqueous solutions: at 300 K, at an elevated temperature of 338 K (65 ℃), and at 338 K coupled with 25 bar pressure. Snapshots at different times for each simulation were examined (Figure 5 ***B***).

The 338 K thermal condition represents the measured temperature that appears upon exposing the aqueous protein solution to sound energy with a power of 2 W, and 20% amplitude. Furthermore, the 338 K temperature value also corresponds to the temperature at which lysozyme fibrillates according to the “classical” protocol for preparing protein fibrils (17, 32–34), whereas the lysozyme protein molecule itself is relatively stable at 338 K, due to its high melting point, varying from ca. 342 K to 347 K at different pH values (35, 36).

Former MD simulations confirmed that relatively high temperatures (370 K) were required to promote and maintain the unfolding process of lysozyme (37). Thus, we hypothesized that at 338 K, the initial stages of structural changes, namely, structural reorganization might take place. To test our hypothesis, the 25-bar (2.5 MPa) pressure condition was chosen, based on the approximate calculation relevant to our experiment, as follows: The effective pressure was estimated from the acoustic power type (ca. 2 W·mm^-2^) and the acoustic impedance of the aqueous solutions (38–40). Considering the density of water at 298-338 K, the speed of sound in water under the same conditions, as well as its density (whose product gives the acoustic impedance), one can estimate that the effective pressure lies in the range of 2.4-2.5 MPa (and of course it increases when the acoustic intensity or the sonication power increases). Considering the presence of the protein in solution and the deviations in ultrasound power, we can conditionally assume an effective pressure of 25 bar (2.5 MPa).

One of the main indicators of the structural stability of proteins during MD simulations is the variability of the root mean square deviation (RMSD) of the C_α_ atoms (*SI Appendix, **Fig. S4***) over time. The changes in RMSD are manifested by the mobility of the protein backbone and the stability of the tertiary structure. Over two runs in an aqueous lysozyme solution at 300 K, the protein exhibited an average C_α_ RMSD value of ∼0.152 ± 0.027 nm with maxima at 0.285 nm. At 338 K, the measured RMSD increased to ∼0.245 ± 0.059 nm, with maxima at 0.408 nm. We observed a similar effect when elevated temperature and pressure were combined, with the mean RMSD increasing to 0.217 ± 0.059 nm, with maxima at 0.445 nm (see **Table 1**).

**Table 1.**
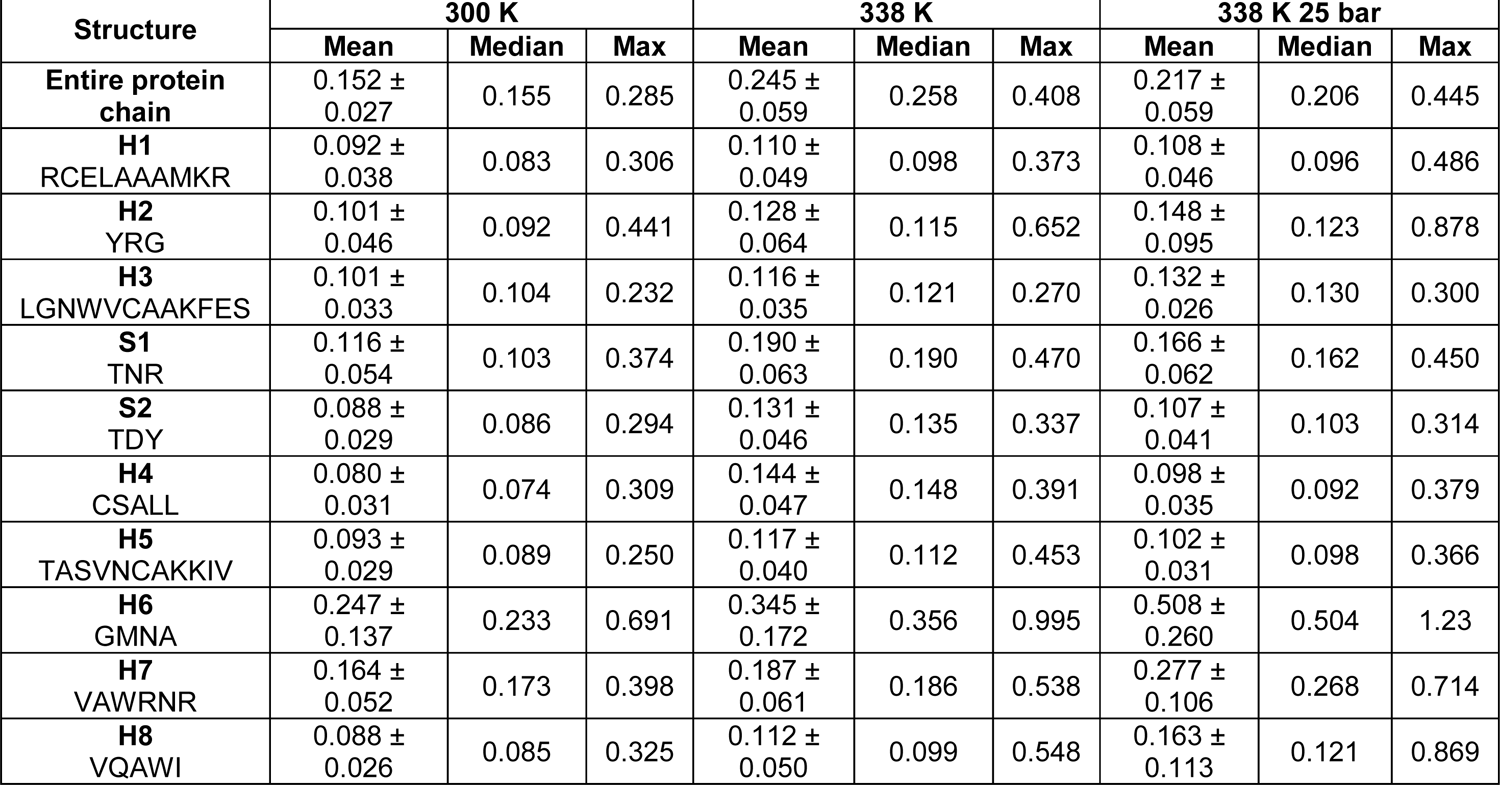
Mean, median, and maximum RMSD value (nm) for C_α_ under different conditions over two runs for 3 µs (with a step of 1 ps) for whole protein chains and each structural element of its secondary structure (referred to in Figure 5 **A**). A graphical representation of the RMSD summary is presented in the SI Appendix, **Fig. S16**.

From the analysis presented in the *SI Appendix, **Fig. S5***, the RMSD value of the C_α_ atoms increases with an increase in either temperature or pressure, indicating protein fold destabilization. At room temperature (RT), namely, at 300 K, our RMSD analysis confirmed that the protein structure is indeed stable (see the *SI Appendix, **Fig. S5***). At elevated temperatures, the RMSD increased to ca. 0.3 nm over a period of approximately 500-750 ns, indicative of partial destabilization of the protein fold. Interestingly, the application of pressure alone (25 bar) stabilizes the protein fold overall, based on RMSD analysis (*SI Appendix, **Fig. S16*** and ***Fig. S18 A***). The combined action of elevated temperature (to 338 K) and pressure (25 bar) enabled us to perform a detailed analysis of the protein fragments and to track the conformational changes in these fragments. The exact sequences of the fragments, which were mentioned earlier (also see Figure 5 ***A***), are summarized in **Table 1**.

Despite the apparent overall stabilization at high pressure (Figure 5 ***C***), the effect on individual structural elements may be more significant (Figure 5 ***D)***. Overall, the simulations carried out revealed changes in all structural elements in proteins, both at elevated temperatures alone and after applying additional pressure. The pressure factor indeed destabilizes the structure, compared with non-elevated temperature conditions (see the *SI Appendix, **Table S1**, **Fig. S16***, and ***Fig. S17***). According to the mean RMSD value, temperature and pressure have a minimal impact on helix H1 (see **Table 1** and the *SI Appendix, **Table S1, Fig. S6**, **Fig. S16**, **Fig. S17**, **Fig. S18***), where C_α_ in H1 did not show any significant changes over time (see the *SI Appendix, **Fig. S6***, and ***Fig. S7***).

The combined effect of temperature and pressure destabilizes the H2 and H3 motifs compared to the temperature alone, concluded from the analysis of RMSD behavior over time (*SI Appendix, **Fig. S7***, and ***Fig. S8***). Furthermore, comparing the RMSD absolute and relative values (**Table 1** and the *SI Appendix, **Table S1***, ***Fig. S16***, and ***Fig. S17***,) as well as the cumulative distribution (*SI Appendix, **Fig. S18***) points to a less significant destabilization effect compared with H6–H8 (see the discussion below).

For S1, S2, H4, and H5, we observed that pressure stabilizes the structure of the fragments, which is opposite to the effect of elevated temperature alone (see **Table 1** and the *SI Appendix, **Table S1**, **Fig. S9**, **Fig. S10**, **Fig. S11**, **Fig. S12***, ***Fig. S16****, **Fig. S17**,* and ***Fig. S18***), with no appreciable trend observed for the RMSD of C_α_ over time.

For fragments H6-H8 (Figure 5 ***D***, see also **Table 1** and the *SI Appendix, **Table S1**, **Fig. S16**, **Fig. S17**,* and ***Fig. S18***), the combined effect of pressure and temperature had a significant destabilization effect on the fragment fold, compared to high temperature alone, which is reflected in the increased values of the mean RMSD. When the temperature and pressure factors were combined, the fold-modifying effect becomes even more pronounced, altering the fragment fold integrity. The dependence on the RMSD of the C_α_ atoms displays a sustainable increasing trend according to the time under high pressure (*SI Appendix, **Fig. S13**, **Fig. S14**,* and ***Fig. S15***), particularly for H6-H8. The H6 fragment was found to be highly sensitive, specifically to thermal fluctuations over time.

Thus, even at temperatures slightly above the RT, variations in RMSD were observed, which became more pronounced over time. Although the conformation of H6-H8 changes, it is essential to note that they do not fully unfold/refold during the 3 µs MD simulations. A more detailed description of MD analysis and discussion is presented in the *SI appendix*). Thus, even though lysozyme does not undergo significant changes in total protein fold, under the investigated conditions, both elevated temperature alone and its combined action with the increased pressure indeed affect certain fragments in the protein structure.

The fragments of H6–H8 were identified by using MD, as structurally sensitive motifs, which are more prone to refolding than the others. Thus, under elevated temperature and pressure, conditions equivalent to those created by exposure to US, fragments undergo partial protein refolding, an essential early step in amyloidogenic protein fibrillation.

An additional, more detailed investigation of the acoustic streaming effects in the context of protein folding could potentially benefit from the microgravity conditions, since they minimize convective currents, thereby isolating the acoustic phenomena. This method would allow a clearer understanding of how ultrasound affects protein aggregation without the interference of gravity-driven convection.

## Conclusions

Our investigation of the effects of ultrasound on the self-assembly of amyloidogenic proteins has yielded significant insights. Our findings indicate that using low-amplitude ultrasound to apply mechanical energy to aqueous media can give rise to relatively mild pressure and temperature variations, which are, however, sufficient to compromise protein complexes’ stability, folding, and self-assembling patterns. Prolonged exposure to low-amplitude ultrasound enables the controlled initiation and further elongation of amyloid protein nanofibrils directly from monomeric lysozyme protein, without introducing pre-formed seeds. Interestingly, the US-grown protein fibrils are characterized by a critical length. Mechanistically, low ultrasonic power levels induce refolding of specific sensitive protein motifs. Molecular dynamics revealed that this refolding originates from pressure changes and is further amplified by temperature variations. Solution X-ray scattering analysis and secondary structure evaluation by FTIR spectroscopy confirmed that the nanofibrillar assemblies created either by ultrasound energy or natively fibrillated lysozyme are structurally identical. In essence, our findings enhance our current understanding of the effect of ultrasound on fibrillar protein self-assembly and highlight its potential applicability in protein chemistry settings.

## Materials and Methods

### Preparation of a lysozyme monomer solution

Monomer solution: Lyophilized hen egg white lysozyme (HEWL, Sigma-Aldrich, USA) was dissolved in 20 mM NaCl + 10 mM HCl at 20 mg·ml^-1^ (1.4 mM). All analyses presented in this paper were conducted using this concentration.

For additional experiments, lysozyme monomer solutions were prepared at various concentrations: 1, 5, 10, 40 mg·ml^-1^ to evaluate the impact of concentration on fibril formation under ultrasound.

### Preparation of lysozyme fibrillar seeds (standard protocol)

The monomer solution of HEWL was incubated for 72 h at 65 ℃. After incubation, a gel was obtained and further sonicated for 4 min by using an Ultrasonic Processor (QSONICA) operating at 20 kHz.

### Formation of lysozyme fibrils under the action of ultrasound

Lysozyme monomer solutions were subjected to sonication using an Ultrasonic Processor (QSONICA) operating at 20 kHz. The sonication was performed at 20% amplitude, corresponding to a delivered power of 2 W, for 10 minutes, after which the resulting fibrils were formed and then analyzed.

The sonication was carried out at 20% amplitude (2 W of delivered power). In sonication processes, power is quantified in watts, whereas amplitude refers to the range of the probe’s tip displacement. The ultrasonic processor ensures a stable amplitude throughout the operation. The intensity of the ultrasonic waves is directly proportional to the amplitude produced by the radiating surface of the tip or horn. When discussing low-power ultrasound in our research, we are considering scenarios where the sonication power is 2 W and the amplitude is set to a relatively low level of 20%. The accumulated energy, measured in Joules and delivered to the probe, is displayed on the instrument for one cycle; it is dependent on the preset amplitude and the exposure duration. In the current research, with an amplitude set at 20% and an exposure time of 10 minutes, the conditions correspond to 1000 Joules.

### Morphological analysis by Atomic Force Microscopy (AFM)

A 50 µl aliquot of the nanofibril solution, formed because of US exposure (2 W, 10 min), was applied to freshly cleaved mica. After 2 minutes, it was thoroughly rinsed with deionized water and air-dried. AFM tapping-mode evaluations were conducted using AC240 probes on a NanoWizard 4 device within a soundproof chamber (JPK Instruments). The images were analyzed using JPK’s Analysis Software, and ImageJ for statistical data extraction was used.

### Field Emission High Resolution Scanning Electron Microscopy (FE-HRSEM)

FE-HRSEM images were obtained using Ultra-55 Ultra-high-resolution SEM (Carl Zeiss, Germany) with an accelerating voltage of 2-3 kV for both In-Lens and Everhart-Thorney secondary electron (SE2) detectors under a vacuum of <2·10^-5^ mbar with a working distance of 2.4-4 mm and an aperture of 30 µm. For imaging, a drop of liquid samples of lysozyme fibrils was spread onto a 2 × 2 cm^2^ mica sheet attached to a 20 × 75 mm^2^ microscope glass for 2 min, gently rinsed with Milli-Q triple distilled water, and dried under a gentle nitrogen stream (like sample preparation in the “Morphological analysis by AFM” section). Before imaging, the slides were attached to aluminum stubs with carbon tape (2SPI, USA), and partially coated with carbon paste (2SPI, USA); finally, 2–3 nm of Iridium was sputtered on the samples using a CCU-010 HV high-vacuum (8·10^-3^ – 5·10^-5^ mbar) sputter coater (Safematic, Switzerland) at a rate of 0.01–0.05 Å·s^-1^ to improve the image contrast, sample conductivity, and stability during imaging.

### AFM nanomechanical measurements

PeakForce Quantitative Nanomechanical Mapping (QNM) has been performed on a Multimode Atomic Force Microscope (AFM) by Bruker and was utilized to capture imaging and mechanical characteristics. Olympus AC160 probes were employed in these measurements, with a known spring constant of 40 N/m. The probe’s effective tip radius, calibrated by using Highly Ordered Pyrolytic Graphite (HOPG) and characterized by an 18 GPa modulus, ensuring deformation depths remained at approximately 1 nm. Elastic modulus mapping was generated via the Nanoscope 9.2 software by Bruker. The Derjaguin-Muller-Toporov (DMT) model was applied to align the force-deformation response curves. Gwyddion, which is open-source software, was engaged for image analysis. The computed average modulus values were extracted from the central region of each fibril.

### Numerical Simulations and Modeling

All numerical analyses, including distribution dynamics, mean-length dynamics, and the comparison of transient distribution with experimental data, were conducted using Wolfram Mathematica version 13.3.1.0.

### Small-angle X-ray scattering (SAXS) measurements

Solution SAXS measurements were performed at ID02 beamline at the European Synchrotron Radiation Facility (ESRF). The beam size was 32.4 x 145 μm^2^ (vertical and horizontal, respectively). The photon energy was 12.23 keV. The detector was an Eiger2 4M (Dectris AG). The sample-to-detector distance was 3.114 m, and the exposure time was 0.1 s (41).

### SAXS Models

All the models were placed so that at their center of mass is at the origin. The models were computed in D+ software with taking into account a Gaussian resolution function with a standard deviation of 0.001, as previously described (24). In addition, we used the following D+ software parameters: Orientation average was computed by Monte Carlo (Mersenne Twister) using 10^6^ Integration iterations. The Grid Size of the monomer was 20 and 300 points were generated, required Convergence of 0.001, using a 500 ms Update interval. We assumed that the protein was hydrated by a 0.28 nm of a solvent layer with an electron density of 340 *e* ⋅ *mm*^-3^. The solvation layer was found using a spherical proble with a radius of 0.14 nm and a solvent voxel size of 0.2 nm. The contribution of the solvent displaced by the protein we used the dummy atom method. The excluded volume parameter of the solvent, *C*_1_, was set to 1, and the solvent electron density was 333 *e* ⋅ *mm*^-3^. The largest reciprocal grid was computed for the monomer and then The Hybrid method was applied for larger structures. These parameters were explained in greater details in our earlier publication (23).

Atomic models of the fibrils were generated by docking copies of the hydrated lysozyme atomic model monomer into the fibril assembly symmetry, containing the rotations and translations of each subunit. The position of a specific monomer, ř_ijk_ = (*x*_i_, *y*_j_, *z*_k_), can be expressed as *x*_i_ = *iD*_x_, *y*_j_ = *jD*_y_, *z*_k_ = *kD*_z_, where *i, j*, and *k* vary within the range of 0 to 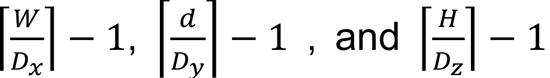, respectively. The values of the fibril height, *H*, depth, *d*, and width, *W*, varied between different fibril models. The spacing *D*_x_, *D*_y_, and *D*_z_ between the centers of adjacent monomers in the *x, y*, and *z* directions, were 3.0 ± 0.1 *nm* in all the models. The Tait-Brain orientation α_ijk_ = β_ijk_ = γ_ijk_= 0 of all the monomers remained unchanged in all the samples. Thermal fluctuations were taken into account by Monte Carlo simulations using a harmonic potential between nearest-neighbor monomers, as explained (27). We then deleted monomers with indexes 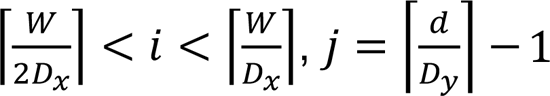 1, and 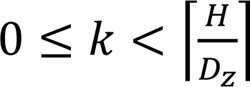, which covered half of the last layer. To optimize the fit of all the models to the scattering data, a very small constant background factor was added.

### Kinetic Analysis Using the ThT assay

To monitor the progression rate of protein fibril formation, we used ThT (Thioflavin T, Sigma-Aldrich, USA) dye, which has an affinity for the β-sheet conformation in protein aggregates. When ThT binds to these structures, its fluorescence intensity markedly increases and red-shifts, making it an exceptionally sensitive indicator for amyloid formations. For the kinetic study, we mixed lysozyme monomer/nanofibril samples with a 20 µM ThT solution. This mixture was then transferred into a 96-well plate (sourced from Greiner Bio-One GmbH, Germany) and incubated at 65 ℃ within a CLARIOstar (BMG LABTECH) plate reader. The ThT fluorescence was captured at excitation and emission wavelengths of 440 nm and 490 nm, respectively. The experiment was concluded once the kinetic saturation phase was observed.

### Protein secondary structure elucidation using FTIR spectroscopy

The measurements were performed in deuterated solvent (D_2_O). A Nicolet iS50 FTIR spectrometer, equipped with an Attenuated Total Reflection (ATR) module from Thermo Scientific, was used for the structural examination of lysozyme fibrils and monomers. The samples were placed directly into the ATR apparatus without additional sample preprocessing. The analysis was performed by initially eliminating the water spectrum from the measurements. Atmospheric noise was subtracted out following the initial collection of FTIR spectra. A second-derivative transformation was applied for more detailed spectral analysis. Finally, the spectral data were processed and interpreted using Thermo Scientific’s OMNIC software and OriginLab’s analytical tools. FTIR band assignment: 1605-1637 cm^-1^ – *β*-sheet content, 1638-1649 cm^-1^ – random coils/extended chains, 1650-1664 cm^-1^ – *α*-helices, 1665-1689 cm^-1^ – *β*-turns, 1690-1705 cm^-1^ – intermolecular *β*-sheets.

### Molecular dynamics simulations

The starting model for hen egg white lysozyme (HEWL) was taken from the PDB databank (PDB ID: 193L). Before simulations, all water in the model was removed; however, the bound sodium and chloride ions were preserved. To somewhat simulate the experimental conditions of pH 2.0, all histidines and carboxylic acid groups were protonated. Gromacs 2022.1 was used with the Amber99SB-ILDN force field and the SPCE water model. The salt concentration was set to 20 mM (NaCl) and somewhat adjusted to neutralize the charge of the system. Following minimization of the solvated system and equilibration with the NVT regime, three separate NPT regimes were carried out for each of the three conditions: room temperature with ambient pressure (300 K), high temperature (338 K) with ambient pressure, and high temperature with high pressure (338 K, 25 bar). This was followed by two production runs under each condition, for 3,000 ns for each run. The molecule was solvated in a water dodecahedron box. The temperature was controlled via a Berendsen temperature coupling thermostat, whereas the pressure was controlled via a Parrinello-Rahman pressure coupling barostat.

The data from the time-dependent RMSD were transformed into the cumulation diagrams presented in *SI Appendix*, ***Figures S5-S15***, and then extracted using Matlab scripts. Structural alignment for presentations and analysis was done in ChimeraX 1.7 using the “matchmaker” command (see Figure 5 ***B***) and in Pymol 1.7/2.5.5 using the “super” and “align” commands.

Coloring by displacement presented in *SI Appendix*, ***Fig. S19***, was done in Pymol using “modevectors” script, coloring by RMSD on the same image was done using “colorbyrmsd” script, whereas coloring by displacement, specifically for the side chain on this figure was done using “colorbydisplacement” script. Pairwise RMSD per residue analysis, similar to the root mean square fluctuation (RMSF), was done using the “rmsdbyres” script.

The values of the *φ* (phi) and *ψ* (psi) angles of the simulated structures for the Ramachandran plots were obtained from Pymol 2.5.5 using the “DynoPlot” script and also in Chimera 1.18. Contour map data for the favored and allowed regions of the angles were generated with Chimera 1.18.

## Acknowledgments

The authors acknowledge the European Synchrotron Radiation Facility (ESRF), beamline ID02 (T. Narayanan, L. Matthews, and the rest of the ID02 team) for providing use of the synchrotron radiation facility and for assistance in using the beamline. U.S. acknowledges financial support from the Gruber Foundation, the Nella and Leon Benoziyo Center for Neurological Diseases. In addition, U.S. thanks the Perlman family for funding the Shimanovich Lab at the Weizmann Institute of Science: “This research was made possible in part by the generosity of the Harold Perlman Family.” The authors would like to acknowledge partial support from the GMJ Schmidt Minerva Center of Supramolecular Architectures at the Weizmann Institute, the Mondry Family Fund for the University of Michigan/Weizmann collaboration, the Gerald Schwartz and Heather Reisman Foundation, and the WIS Sustainability and Energy Research Initiative (SAERI). This research was supported by a research grant from the Tom and Mary Beck Center for Advanced and Intelligent Materials at the Weizmann Institute of Science, Rehovot, Israel. Molecular graphics and analyses were performed with UCSF ChimeraX, developed by the Resource for Biocomputing, Visualization, and Informatics at the University of California, San Francisco, with support from National Institutes of Health R01-GM129325 and the Office of Cyber Infrastructure and Computational Biology, National Institute of Allergy and Infectious Diseases. A.A. thanks the Clore center for Biological Physics for their support. The authors are also grateful to Steve Manch for the English editing.

## References

1. T. P. J. Knowles, M. J. Buehler, Nanomechanics of functional and pathological amyloid materials. Nat. Nanotechnol. 6, 469–479 (2011).

2. F. Chiti, C. M. Dobson, Protein misfolding, amyloid formation, and human disease: A summary of progress over the last decade. Annu. Rev. Biochem. 86, 27–68 (2017).

3. M. B. Pepys, et al., Human lysozyme gene mutations cause hereditary systemic amyloidosis. Nature 362, 553–557 (1993).

4. L. Nielsen, et al., Effect of environmental factors on the kinetics of insulin fibril formation: Elucidation of the molecular mechanism. Biochemistry 40, 6036–6046 (2001).

5. R. Gallardo, N. A. Ranson, S. E. Radford, Amyloid structures: much more than just a cross-β fold. Curr. Opin. Struct. Biol. 60, 7–16 (2020).

6. X. Periole, A. Rampioni, M. Vendruscolo, A. E. Mark, Factors that affect the degree of twist in β-sheet structures: A Molecular dynamics simulation study of a cross-β filament of the GNNQQNY peptide. J. Phys. Chem. B 113, 1728–1737 (2009).

7. L. Lu, Y. Deng, X. Li, H. Li, G. E. Karniadakis, Understanding the Twisted Structure of Amyloid Fibrils via Molecular Simulations. J. Phys. Chem. B 122, 11302–11310 (2018).

8. T. P. J. Knowles, et al., An analytical solution to the kinetics of breakable filament assembly. Science (80-.). 326, 1533–1537 (2009).

9. A. Weissler, I. Pecht, M. Anbar, Ultrasound chemical effects on pure organic liquids. Science (80-.). 150, 1288–1289 (1965).

10. C. L. Hawkins, M. J. Davies, Generation and propagation of radical reactions on proteins. Biochim. Biophys. Acta - Bioenerg. 1504, 196–219 (2001).

11. U. Shimanovich, et al., Sonochemically-induced spectral shift as a probe of green fluorescent protein release from nano capsules. RSC Adv. 4, 10303–10309 (2014).

12. U. Shimanovich, et al., Tetracycline nanoparticles as antibacterial and gene-silencing agents. Adv. Healthc. Mater. 4, 723–728 (2015).

13. U. Shimanovich, G. J. L. Bernardes, T. P. J. Knowles, A. Cavaco-Paulo, Protein micro- and nano-capsules for biomedical applications. Chem. Soc. Rev. 43, 1361–1371 (2014).

14. O. Grinberg, U. Shimanovich, A. Gedanken, Encapsulating bioactive materials in sonochemically produced micro- and nano-spheres. J. Mater. Chem. B 1, 595–605 (2013).

15. F. Grigolato, P. Arosio, Synergistic effects of flow and interfaces on antibody aggregation. Biotechnol. Bioeng. 117, 417–428 (2020).

16. A. Kozell, A. Solomonov, U. Shimanovich, Effects of sound energy on proteins and their complexes. FEBS Lett. 597, 3013–3037 (2023).

17. A. Kozell, et al., Modulating amyloids’ formation path with sound energy. Proc. Natl. Acad. Sci. U. S. A. 120, e2212849120 (2023).

18. W. H. Wu, P. F. Yang, W. Zhai, B. B. Wei, Oscillation and Migration of Bubbles within Ultrasonic Field. Chinese Phys. Lett. 36, 084302 (2019).

19. M. Ehsani, N. Zhu, H. Doan, A. Lohi, A. Abdelrasoul, In-situ synchrotron X-ray imaging of ultrasound (US)-generated bubbles: Influence of US frequency on microbubble cavitation for membrane fouling remediation. Ultrason. Sonochem. 77, 105697 (2021).

20. L. A. Morozova-Roche, et al., Amyloid fibril formation and seeding by wild-type human lysozyme and its disease-related mutational variants. J. Struct. Biol. 130, 339–351 (2000).

21. L. N. Arnaudov, R. De Vries, Thermally induced fibrillar aggregation of hen egg white lysozyme. Biophys. J. 88, 515–526 (2005).

22. N. P. Reynolds, et al., Competition between crystal and fibril formation in molecular mutations of amyloidogenic peptides. Nat. Commun. 8, 1338 (2017).

23. A. Ginsburg, et al., D+: software for high-resolution hierarchical modeling of solution X-ray scattering from complex structures. J. Appl. Crystallogr. 52, 219–242 (2019).

24. E. Balken, et al., Upgrade of D+ software for hierarchical modeling of X-ray scattering data from complex structures in solution, fibers and single orientations. J. Appl. Crystallogr. 56, 1295–1303 (2023).

25. T. D. Goddard, et al., UCSF ChimeraX: Meeting modern challenges in visualization and analysis. Protein Sci. 27, 14–25 (2018).

26. E. F. Pettersen, et al., UCSF ChimeraX: Structure visualization for researchers, educators, and developers. Protein Sci. 30, 70–82 (2021).

27. D. Louzon, et al., Structure and Intermolecular Interactions between L-Type Straight Flagellar Filaments. Biophys. J. 112, 2184–2195 (2017).

28. S. A. Khrapak, Lindemann melting criterion in two dimensions. Phys. Rev. Res. 2 (2020).

29. H. Naiki, K. Higuchi, M. Hosokawa, T. Takeda, Fluorometric determination of amyloid fibrils in vitro using the fluorescent dye, thioflavine T. Anal. Biochem. 177, 244–249 (1989).

30. M. Biancalana, K. Makabe, A. Koide, S. Koide, Molecular Mechanism of Thioflavin-T Binding to the Surface of β-Rich Peptide Self-Assemblies. J. Mol. Biol. 385, 1052–1063 (2009).

31. L. W. Y. Roode, U. Shimanovich, S. Wu, S. Perrett, T. P. J. Knowles, Protein Microgels from Amyloid Fibril Networks. Adv. Exp. Med. Biol. 1174, 223–263 (2019).

32. M. Xu, et al., The first step of hen egg white lysozyme fibrillation, irreversible partial unfolding, is a two-state transition. Protein Sci. 16, 815–832 (2007).

33. X. M. Zhou, et al., Enzymatically Active Microgels from Self-Assembling Protein Nanofibrils for Microflow Chemistry. ACS Nano 9, 5772–5781 (2015).

34. U. Shimanovich, et al., Protein microgels from amyloid fibril networks. ACS Nano 9, 43–51 (2015).

35. S. Venkataramani, J. Truntzer, D. R. Coleman, Thermal stability of high concentration lysozyme across varying pH: A Fourier Transform Infrared study. J. Pharm. Bioallied Sci. 5, 148–153 (2013).

36. C. O. Silva, et al., Lysozyme photochemistry as a function of temperature. the protective effect of nanoparticles on lysozyme photostability. PLoS One 10, e0144454 (2015).

37. M. Jafari, F. Mehrnejad, Molecular insight into human lysozyme and its ability to form amyloid fibrils in high concentrations of sodium dodecyl sulfate: A view from molecular dynamics simulations. PLoS One 11, e0165213 (2016).

38. B. Sorum, R. A. Rietmeijer, K. Gopakumar, H. Adesnik, S. G. Brohawn, Ultrasound activates mechanosensitive TRAAK K+ channels through the lipid membrane. Proc. Natl. Acad. Sci. U. S. A. 118, e2006980118 (2021).

39. R. Hatakeyama, M. Yoshizawa, T. Moriya, Method for the measurement of acoustic impedance and speed of sound in a small region of bone using a fused quartz rod as a transmission line. Japanese J. Appl. Physics, Part 1 Regul. Pap. Short Notes Rev. Pap. 39, 6449–6454 (2000).

40. R. Alkins, K. Hynynen, Ultrasound Therapy. Compr. Biomed. Phys. 10, 153–168 (2014).

41. T. Narayanan, et al., Performance of the time-resolved ultra-small-angle X-ray scattering beamline with the Extremely Brilliant Source. J. Appl. Crystallogr. 55, 98–111 (2022).

