## Supplementary information for "Sound-mediated nucleation and growth of amyloid fibrils"

**Supporting Information Text 1**

**Model for dynamics of the fibril length distribution**

**Model formulation**

This section presents a simplified model to describe the dynamics observed in our experimental setup. Previous works have developed similar kinetic models, following the pioneering work of Ref. (1). The model we introduced here is a particular case of the general model studied in Refs. (2-4), which enabled us to obtain analytical results for the steady-state distributions. Although these results are partially known from Ref. (4), we included them in the text for completeness. For a more extensive reference list on the kinetic approach to describe amyloid fibrils, see a recent review (5).

We consider fibrils as linear chains of elementary building blocks (monomers), and measure polymer length, $x$, in discrete units corresponding to the number of monomers. First, we consider a model in which the local dynamics are governed by three fundamental processes: dimerization (the nucleation of a new fibril from two monomers), elongation of an existing fibril, and fragmentation of an existing fibril into two fibrils. We will see that the steady state of this model cannot reproduce the experimental results – for any chosen parameter– however, its transient behavior can match the measurements well. In the last section, we will extend the model to include an additional process, the detachment of a single monomer from the tip of the fibril, and we will show that this alternative model can also reproduce the experimentally observed long fibrils also in steady state. Finally, we will discuss future means for distinguishing between the two models.

Despite the complexity of the non-equilibrium kinetics induced by the ultrasound, we consider a coarse-grained model (hereafter referred to as the “basic model”) wherein the rate of elongation $\alpha n_{1}$ is proportional to the concentration of free monomers (i.e. we assume first-order kinetics) and fragmentation of a fibril of length $x$ can occur with a constant rate $\nu$ at each of the $x-1$ interfaces between monomers. Similarly, dimerization is assumed to proceed at a rate $\alpha_{1}n_{1}^{2}$ proportional to the square of the free monomer concentration, since it involves the interaction of two monomers. The model is schematically presented in ***Figure 2 A (1)-(3)*** in the Main Text***.***

Our primary interest lies in tracking the concentration of fibrils $n_{x}(t)$ with a specific length $x$ over time (i.e., $n_{1}(t)$ corresponds to the free monomer concentration).

We begin with finding the steady-state distribution of fibril length. To this end, we formulate the master equation, which describes the temporal evolution of the concentrations:

$$\begin{matrix} \frac{dn_{1}}{\text{ }dt} & =-2\alpha_{1}n_{1}^{2}+2\nu\sum_{y=2}^{\infty} n_{y}(t)-\alpha n_{1}\sum_{y=2}^{\infty} n_{y}, \\ \frac{dn_{2}}{\text{ }dt} & =\alpha_{1}n_{1}^{2}-\alpha n_{1}n_{2}-\nu n_{2}+2\nu\sum_{y=3}^{\infty} n_{y}(t), \\ \frac{dn_{x}}{\text{ }dt} & =-\alpha n_{1}\left( n_{x}-n_{x-1} \right)-\nu(x-1)n_{x}+2\nu\sum_{y=x+1}^{\infty} n_{y}(t), x>2. \end{matrix}$$

*( 1 )*

The factor of 2 before $\alpha_{1}$ in the first equation accounts for both monomers vanishing during dimerization. The factor of 2 before $\nu$ acknowledges the two ways a fibril of length $x$ can break into fragments of length $y$ and $x-y$, except for $x=2y$, where there is only one choice for the breaking site, but both fragments contribute to $n_{y}$.

The master equation can be rewritten succinctly as:

$$\begin{matrix} & \frac{dn_{x}}{\text{ }dt}=-\left( \alpha-\alpha_{1} \right)n_{1}\left( \delta_{x,2}n_{x-1}-\delta_{x,1}n_{x} \right)-\alpha n_{1}\left( n_{x}-\left( 1-\delta_{x,1} \right)n_{x-1} \right)- \\ & -\nu(x-1)n_{x}+2\nu\sum_{y=x+1}^{\infty} n_{y}(t)+\delta_{x,1}\left( \alpha-\alpha_{1} \right)n_{1}^{2}-\delta_{x,1}\alpha n_{1}\sum_{y=1}^{\infty} n_{y}. \end{matrix}$$

*( 2 )*

It is easy to check that this equation conserves the total concentration of monomers (either free or in fibrils) $\mathcal{N}\overset{\text{ def }}{=}\sum_{x=1}^{\infty} xn_{x}$, but note that $\sum_{x=1}^{\infty} n_{x}$ is not conserved. Hereafter, we will assume that the initial condition consists solely of monomers and no fibrils, i.e. $n_{x}(0)=\mathcal{N}\delta_{1,x}$.

The above model has 4 parameters: $\alpha,\alpha_{1},\nu$, and $\mathcal{N}$. There are several dimensionless constants that will be important for determining the behavior: $\lambda\overset{\text{ def }}{=}\alpha\mathcal{N}/\nu$, which compares the elongation rate to that of fragmentation (which we will later assume is $\gg1$ ). Similarly, $\lambda_{1}\overset{\text{ def }}{=}\alpha_{1}\mathcal{N}/\nu$ compares the rate of dimerization to fragmentation, and in order to explain the experimental data, we will later assume that $\lambda_{1}\ll\lambda$. Surprisingly, we will show that even for such a small fragmentation rate, at steady-state, the distribution of fibrils does not contain long fibrils (in fact, the mean fibril length is precisely 3 in this limit). Deviating the parameters $\lambda$ and $\lambda_{1}$ from the assumed range would only make the fibril distribution tend to smaller values, hence, our conclusions do not hinge on this assumption.

We will first consider the steady-state (achieved at long times and independent of the initial conditions), and show that it generically leads to a small mean fibril length, inconsistent with the experimental data. We also find that for $\lambda\gg1$ and $\lambda\gg\lambda_{1}$ the steady-state distribution is universal and independent of the model parameters.

This motivated us to consider the dynamics of the length distribution. Using numerical simulations, we show that long fibrils can exist at intermediate times, and the mean fibril length is non-monotonic in time. A comparison of the predicted transient distribution and the experimental data shows good agreement, suggesting that the experiments probe this regime.

**Mean fibril length**

Utilizing the master equation, we can readily formulate a closed set of equations, similar to those proposed by, e.g., (2), (6), (7), for the total fibril length per unit volume, denoted as $\mathcal{l}_{T}\overset{\text{ def }}{=}\sum_{x=2}^{\infty} xn_{x}$, the total concentration of fibrils $n_{T}\overset{\text{ def }}{=}\sum_{x=2}^{\infty} n_{x}$, and the concentration of free monomers $n_{1}$ :

$$\begin{matrix} \frac{dn_{1}}{\text{ }dt}=2\nu n_{T}-2\alpha_{1}n_{1}^{2}-\alpha n_{1}n_{T} \\ \frac{dn_{T}}{\text{ }dt}=\alpha_{1}n_{1}^{2}+\nu\mathcal{l}_{T}-3\nu n_{T} \\ \frac{d\mathcal{l}_{T}}{\text{ }dt}=-2\nu n_{T}+2\alpha_{1}n_{1}^{2}+\alpha n_{1}n_{T} \end{matrix}$$

*( 3 )*

Note that $\dot{n}_{1}+{\dot{\mathcal{l}}}_{T}\equiv\dot{\mathcal{N}}=0$ as it should be (overall number of monomers conserves). Setting ${\dot{\mathcal{l}}}_{T}=0$ and $\dot{n}_{T}=0$ (steady-state) one gets:

$n_{T}=\frac{2\alpha_{1}n_{1}^{2}/\nu}{2-\alpha n_{1}/\nu}, \mathcal{l}_{T}=3n_{T}-\alpha_{1}n_{1}^{2}/\nu, \langle x\rangle\overset{\text{ def }}{=}\frac{\mathcal{l}_{T}}{n_{T}}=2+\frac{\alpha n_{1}/\nu}{2}$,

*( 4 )*

$$\alpha_{1}\alpha n_{1}^{3}/\nu^{2}+\left( 4\alpha_{1}/\nu-\alpha/\nu\right)n_{1}^{2}+(2+\alpha\mathcal{N}/\nu)n_{1}-2\mathcal{N}=0.$$

*( 5 )*

The last equation provides the number of free monomers in a steady-state, which, as observed, is also responsible for the mean fibril length $\langle x\rangle$. Surprisingly, these equations yield $\langle x\rangle\leq3$ for all possible values of the parameters: to see this, note that the expression for $n_{T}$ in Eq.(4) must be non-negative, which automatically implies an upper limit of $n_{1}\leq\frac{2}{\alpha/\nu}$ on the free monomers concentration, and, consequently, an upper limit of $\left\langle x \right\rangle\leq3$ on the mean length.

To proceed, we assume, as mentioned above, that $\lambda\gg1$ and $\lambda\gg\lambda_{1}$, which in combination automatically yields $\delta\overset{\text{ def }}{=}\lambda_{1}/\lambda^{2}\ll1$. Under these assumptions, the solution to Eq. (5) can be approximated as:

$$n_{1}=\frac{\nu}{\alpha}\left[ 2-24\delta+\mathcal{O}(\delta/\lambda)+\mathcal{O}\left( \delta^{2} \right) \right] \Longrightarrow\langle x\rangle=3-12\delta+\mathcal{O}(\delta/\lambda)+\mathcal{O}\left( \delta^{2} \right)$$

*( 6 )*

Therefore, the leading term governing the mean fibril length is 3. Consequently, we observe that the fibrils tend to be very short in a steady-state. It is also worthwhile to note that in this limit, the relative fraction of monomers to fibrils $\left( n_{1}/n_{T} \right)$ is negligible: to see this, note that plugging in the leading order contribution $n_{1}\approx\frac{2}{\alpha/\nu}$ into the expression of $n_{T}$ leads to a divergence. Finally, we note that the mean fibril length only hits the bound of $\langle x\rangle=3$ for $\delta=0$, which means that relaxing the assumptions of $\lambda\gg1$ and $\lambda\gg\lambda_{1}$ will only lead to shorter mean fibril length. In other words, no matter what the model parameters are, it will only contain short fibrils in steady-state, and it will not be able to explain the experimental observation of very long fibrils.

For completeness, we next derive an explicit formula for the steady-state fibril distribution, under the assumptions of $\lambda\gg1$ and $\lambda\gg\lambda_{1}$.

**Steady-state solution**

To find the steady-state distribution, we set the temporal derivative in the master equation (2) to zero and subtract the equations for $n_{x+1}$ and $n_{x}$ to obtain

$$\begin{matrix} & \alpha n_{1}/\nu\left( n_{x-1}\left( 1-\delta_{x,1} \right)-2n_{x}+n_{x+1} \right)-(x-1)n_{x}+ \\ & +xn_{x+1}+2n_{x+1}+\left( 3\delta_{x,1}-\delta_{x,2} \right)\left( \alpha/\nu-\alpha_{1}/\nu\right)n_{1}^{2}-\alpha\delta_{x,1}n_{1}n/\nu=0, \end{matrix}$$

*( 7 )*

with $n=n_{T}+n_{1}$ (i.e. the total number of structures, both monomers and mature fibrils). We will next write an equation for the fibril distribution, characterized by the relative fraction $P_{x}=n_{x}/n$ of a fibril of a given length (including $x=1$ ). To this end, we divide Eq. (7) by $n$, and utilize the fact that for the steady-state under the assumptions of $\lambda\gg1$ and $\lambda\gg\lambda_{1}$, as explained above, $n_{1}/n\ll1$. This leads to:

$$2\left( P_{x-1}\left( 1-\delta_{x,1} \right)-2P_{x}+P_{x+1} \right)-(x-1)P_{x}+xP_{x+1}+2P_{x+1}-2\delta_{x,1}=0$$

*( 8 )*

Substituting $x=1$ one finds that $P_{2}=2/5$ and the equation simplifies to

$$(x+5)P_{x+2}-(x+4)P_{x+1}+2P_{x}=0, P_{1}=0, P_{2}=2/5$$

*( 9 )*

Acting on this equation with $\sum_{x} z^{x}$ yields a differential equation on the generating function $G(z)$, with the solution

$$G(z)\overset{\text{ def }}{=}\sum_{x} z^{x}P_{x}=\frac{2z^{3}+3z^{2}+3e^{2z}(z-1)^{2}-3}{2z^{3}}$$

*( 10 )*

Finally, by Taylor-expanding this generating function, we obtain the closed expression for the probabilities:

$$P_{x}=3\frac{2^{x}(x+2)(x-1)}{(x+3)!}$$

*( 11 )*

It can be readily checked that this distribution is normalized and solves Eq. (9) by direct substitution. As we can see, the steady-state asymptotically leads to a universal distribution with no characteristic scale and mean length equal to 3, which contradicts the experimental data (***Figure 2 B*** in the Main text). Although we assumed $\lambda\gg1$ and $\lambda\gg\lambda_{1}$, deviations from these assumptions will only make the experiment discrepancy more severe.

The origin of the inevitably short fibril distribution is as follows: over time, longer fibrils that may have formed will eventually disintegrate, and if a long fibril undergoes fragmentation when the monomer pool is depleted, there is no mechanism to regain the length for it due to the vanishing concentration of free monomers. These free monomers are only sourced by infrequent detachment events from longer fibrils.

**Transient state and connection to experiments**

As mentioned earlier, the steady-state distribution only concentrates on short fibrils in contrast with the observed data. We therefore investigated whether the model can produce long fibrils at intermediate times (i.e. as a transient phenomenon).

The rationale behind the potential existence of long fibrils during transient states can be summarized as follows: starting from an initial "all monomers" state, if the dimerization rate is significantly larger than the elongation rate, free monomers tend to form dimers rapidly, resulting in a large number of short fibrils. Conversely, when the dimerization rate is lower, after the production of the first dimer nuclei, they promptly attract all available free monomers, facilitating the production of longer fibrils that will fragment afterwards. In such a regime, the mean fibril length will follow a non-monotonic trajectory over time, initially increasing and then decaying to asymptotically reach the steady-state value of $\langle x\rangle=3$.

We indeed observe these non-monotonic dynamics by numerically solving the system of Eqs. (3), as shown in **Fig. S1**.

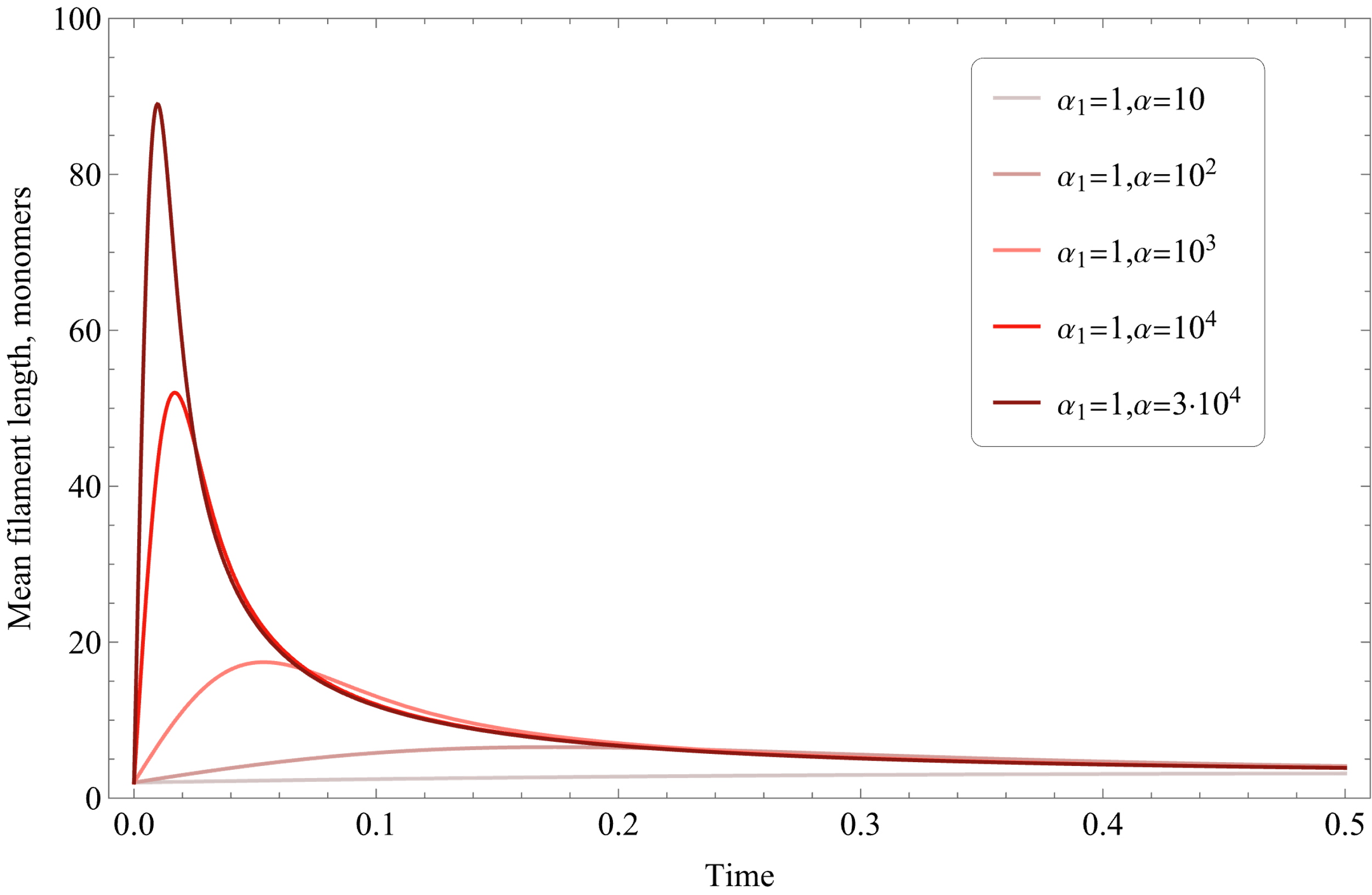

### Fig. S1. The mean fibril length $\boldsymbol{\langle}\boldsymbol{x}\boldsymbol{\rangle}\overset{\text{ def }}{\mathbf{=}}\mathcal{l}_{\boldsymbol{T}}\mathbf{/}\boldsymbol{n}_{\boldsymbol{T}}$ as a function of time numerically computed as a solution for the system (3) for fixed $\boldsymbol{\nu}\mathbf{=1,}\boldsymbol{\alpha}_{\mathbf{1}}\mathbf{=1}$ and different values of $\boldsymbol{\alpha}$ within the time interval $\boldsymbol{t}\boldsymbol{\in[0,0.5]}$. The initial conditions are given by $\boldsymbol{n}_{\mathbf{1}}\mathbf{(0)=1,}\boldsymbol{n}_{\boldsymbol{T}}\mathbf{(0)=0,}\mathcal{l}_{\boldsymbol{T}}\mathbf{(0)=0}$. The calculation is implemented in Wolfram Mathematica NDSolve with the explicit Runge-Kutta method. Non-monotonic behavior and large transient lengths are observed for larger $\boldsymbol{\alpha}$.

Importantly, by choosing a particular set of parameters, the transient dynamics reproduce the experimentally observed fibril length distribution, as shown in ***Figure 2*** **B** in the Main Text. The probability density function (PDF) for the transient state was initialized as $n_{x}(0)=\delta_{1,x}$ and computed with parameters $\nu=1,\alpha_{1}=10,\alpha={10}^{6}$ at time $t=9\cdot{10}^{-3}$.

**Modified model**

So far, we have shown that the basic model reproduces the observed histogram transiently, see ***Figure 2 A and B*** in the Main Text, while the steady-state universally generates only short filaments. We then argued whether there is a steady-state explanation beyond the scope of the basic model.

As mentioned earlier, the limiting factor preventing the master equation (1) from providing long fibrils in a steady state is the insufficient number of free monomers; therefore, to explain the data, one should consider a mechanism that increases the number of free monomers in the steady state. Such a mechanism, namely the *monomer detachment* process, was already present in seminal works by Oosawa (1) and others. We also introduce this process occurring at rate $\beta$ (see ***Figure 2 A (4)*** in the Main Text), so the master equation reads as

$$\begin{matrix} \frac{dn_{1}}{\text{ }dt} & =-2\alpha_{1}n_{1}^{2}+(2\nu+2\beta)\sum_{y=2}^{\infty} n_{y}(t)+2\beta n_{2}-\alpha n_{1}\sum_{y=2}^{\infty} n_{y} \\ \frac{dn_{2}}{\text{ }dt} & =\alpha_{1}n_{1}^{2}-\alpha n_{1}n_{2}-\nu n_{2}+2\nu\sum_{y=3}^{\infty} n_{y}(t)+2\beta n_{3}-2\beta n_{2}, \\ \frac{dn_{x}}{\text{ }dt} & =-\alpha n_{1}\left( n_{x}-n_{x-1} \right)-\nu(x-1)n_{x}+2\nu\sum_{y=x+1}^{\infty} n_{y}(t)+2\beta n_{x+1}-2\beta n_{x}, x>2. \end{matrix}$$

*( 12 )*

The modified model contains one more dimensionless parameter, $\beta/\nu$, which quantifies how frequent the detachment events are in comparison with the fragmentation events. Following the logic explained earlier, we aim for a steady-state solution with a high concentration of free monomers, which implies that an appropriate limit is $\beta/\nu\gg1$.

Solving the system (12) numerically reveals that such a model indeed may reproduce the data in the steady state (see ***Figure 2 B*** in the Main Text). The steady-state PDF for the modified model is computed with parameters $\nu=1, \alpha_{1}={10}^{9}, \alpha=1.2\cdot{10}^{12}, \beta=0.6\cdot{10}^{11}, \mathrm{and} \mathcal{N}=1$.

**Concentration dependency experiment and model validation**

To assess both presented models, we conducted experiments measuring the final mature fibrils’ mass fraction as a function of the initial monomer concentration. Given that both models fit the data in ***Figure 2 B*** in the Main Text with the corresponding solutions initialized as $n_{x}(0)=\delta_{1,x}$, and that this data corresponds to the initial concentration of 20 mg∙ml^–1^, we rescaled the model parameters for the initial condition, $n_{x}(0)=20\frac{mg}{ml}\delta_{1,x}$.

The adjusted parameters are $\nu=1,\alpha_{1}=0.5,\alpha={0.5\cdot10}^{5}$ at time $t=9\cdot{10}^{-3}$ for the basic model and $\nu=1, \alpha_{1}=0.5\cdot{10}^{8}, \alpha=0.6\cdot{10}^{11}, \mathrm{and} \beta=0.6\cdot{10}^{11}$ for the modified model (the execution time for the last one is assumed to be enough to reach the steady state).

We solved both the master equation (1) and the modified master equation (12) with these parameters for various values of the total monomer concentration$\mathcal{N}$ to determine the dependence of the fibril mass fraction $1-n_{1}(\mathcal{N)}/\mathcal{N}$ on $\mathcal{N}$. A comparison of the calculated curves with the measured data revealed that the modified model provides a better fit than the basic one, especially for higher values of the initial monomer concentrations (see ***Figure 2 C*** in the Main Text).

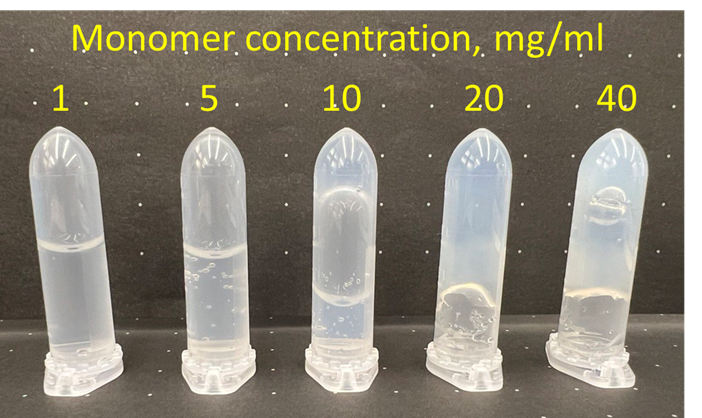

Fig. S2. A photograph illustrating the samples treated by ultrasound, which vary depending on the protein concentration, refers to the *Figure 2 C* in the Main Text.

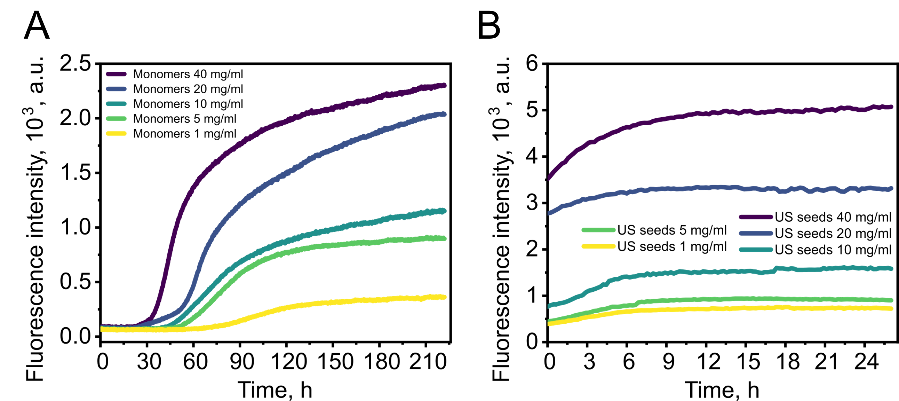

Fig. S3. Non-normalized kinetic curves of data set presented in *Figure 4* *A* in the Main Text.

Supporting Information Text 2

**Molecular dynamics (MD) simulations**

Overall, the simulations show that lysozyme has a stable, weakly varying frame, consisting of S1, S2, H4, and H5, which becomes even more stable under pressure at elevated temperature as well as a highly variable region, consisting of H6-H8, whose variability under pressure gradually decreases with pressure. Regions of H1-H3 are semi-stable, and pressure makes changes in these units but less than in H6-H8.

To display the changes in protein structure, we took a random frame during the simulation at 300 K and overlaid it with the structures at 338 K (*t* = 449693 ps) and two states at 338 K 25 bar displaying changes in H7 in the time frame when RMSD for H7 is 0.35 nm (state 1, *t* = 327309 ps) and when it reaches 0.5 nm (state 2, *t* = 1843799 ps). ***Fig. S19*** shows the pairwise structural alignment of the structures mentioned above and shows the main secondary units, side chain displacement, and regions with high and low RMSD changes. ***Fig. S20***, ***Fig. S21***, and ***Fig. S22*** show the structural alignment of separate elements over the structure at 300 K, displaying maximum changes of the elements in these timeframes. Performed pairwise RMSD per residue analysis (***Fig. S23***, and ***Fig. S24***) for these structures revealed that the random coil may also impact the structural mobility of the protein, and sometimes even more than the ordered units, but do not influence the structural content. It was also noted that the side chain is also very mobile compared with the backbone protein structure.

During the simulations, we found that the secondary structure of the protein undergoes changes regarding the secondary structure composition. The data presented in ***Fig. S25*** indicate that the helices of H1, H3, H6, H7, and H8 tend to become shorter with increasing temperature as well as with additional pressure applied at both states. H2 is the most stable motif, whereas H4 and H5 do not show any tendency to become shortened. However, it was noted that at 338 K 25 bar (state 2) the *β*-sheet structure becomes longer for one amino acid, overall stabilizing the structure.

Secondary structure composition analysis, presented in ***Table S1*** and ***Fig. S26***, showed a general trend of a decrease in the percentage of *α*-helixes and an increase in the percentage of loops and *β*-sheets. The same but a sharper trend was observed by Jafari and Mehrnejad (8) while studying the structural transitions of human lysozyme protein at a high temperature (370 K). However, despite the changes within individual units in the protein structure, the temperature and pressure do not significantly influence the overall structural composition of the protein; however, there are some early stages where the appearance of elongated *β*-sheet structures might be detected.

The Ramachandran plot is one type of protein structure analysis that allows the visualization of the dihedral angles *φ* (phi) and *ψ* (psi) of the protein backbone relative to the central atom of an amino acid C_α_ (***Fig. S27***). Because of steric hindrance between adjacent atoms within a protein structure, the values of these angles are usually limited to certain areas of the graph, especially for helices and sheets. Dihedral angles for loops can occupy any region that is sterically permitted. Owing to the absence of bulky substituents, certain amino acids (*e.g.*, glycine) may have more degrees of freedom to rotate, leading to a rather broad distribution of these angles, whereas others, such as proline, have a more limited distribution. The above-mentioned amino acids represent extreme cases of these two events. On the one hand, the Ramachandran plot is a quite convenient way to represent changes in protein structure during molecular dynamics simulation and to show the distribution of dihedral angles. On the other hand, it is a way to validate the geometry of a protein against the background of preferred and/or allowed regions compiled previously, based on similar calculations/experimental data for many other proteins.

As follows from the Ramachandran plots all the ***Fig. S28*** despite the scatter of RMSD especially for H6-H8, almost none of dihedral angles in the amino acid residues of the protein (with the exception of labile glycine and serine in the case of 338 K 25 bar, state 2) move into regions outside allowed, which indicates the reliability of the calculations and the likelihood of the existence of these structures. However, the distribution in each case differs, evidencing structural rearrangements in the protein molecule.

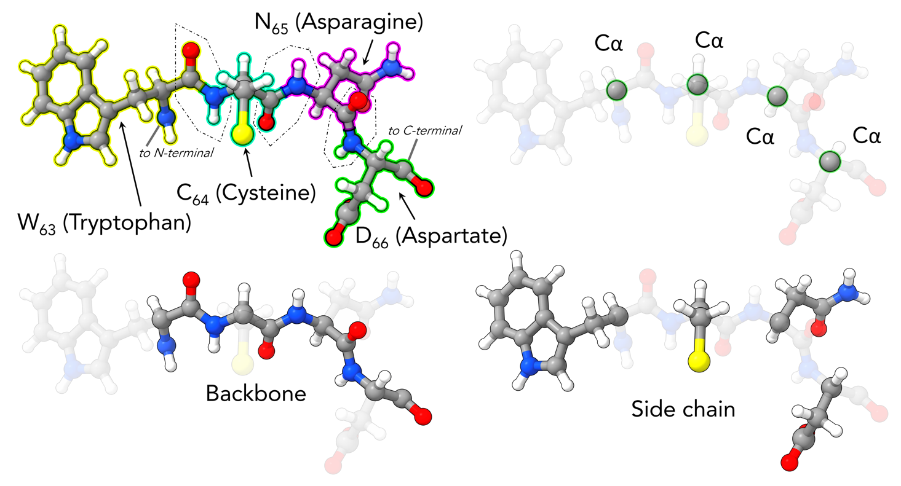

Fig. S4. Top left: a part of the HEWL sequence (63–66, WCND) displaying the main elements of the structure of the amino acid residues and peptide bongs. Top right – carbon alpha (C_α_) positions in the corresponding sequence; bottom left – protein backbone; bottom right – protein side chain elements. Visualization: ChimeraX 1.7.

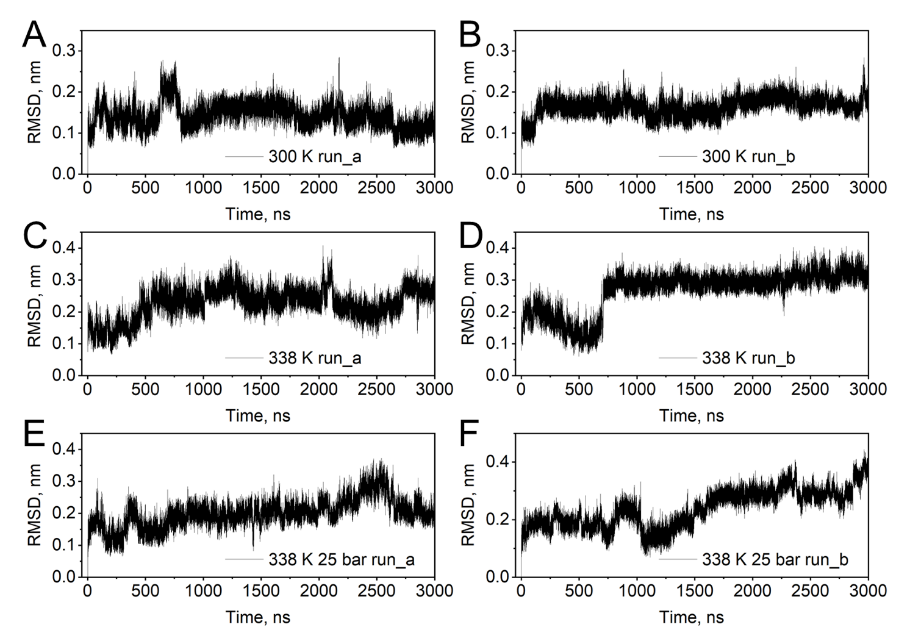

Fig. S5. RMSD of protein C_α_ for residues 1-129 (the entire protein chain) for two independent runs at (a, b) 300 K, (c, d) 338 K, and (e, f) 338 K 25 bar for 3 microseconds.

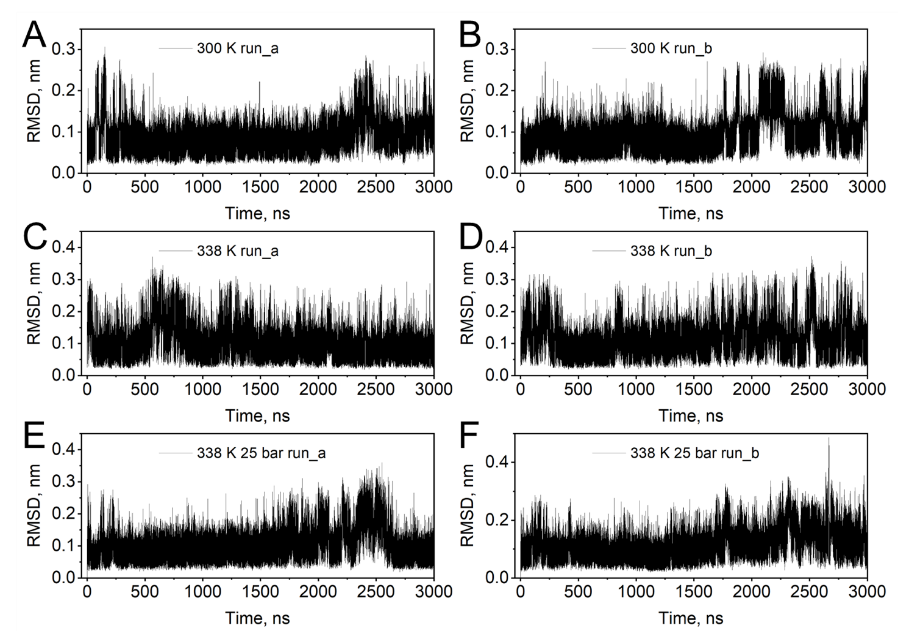

Fig. S6. RMSD of protein C_α_ for residues corresponding to H1 (see *Figure 5 A* in the Main Text) for two independent runs at (a, b) 300 K, (c, d) 338 K, and (e, f) 338 K 25 bar for 3 microseconds.

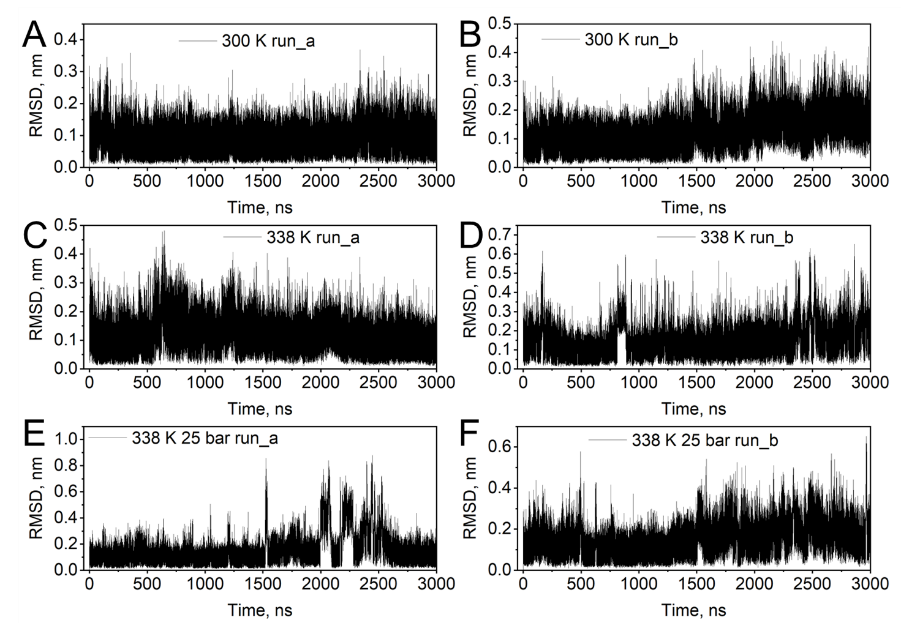

Fig. S7. RMSD of protein C_α_ for residues corresponding to H2 (see *Figure 5 A* in the Main Text) for two independent runs at (a, b) 300 K, (c, d) 338 K, and (e, f) 338 K 25 bar for 3 microseconds.

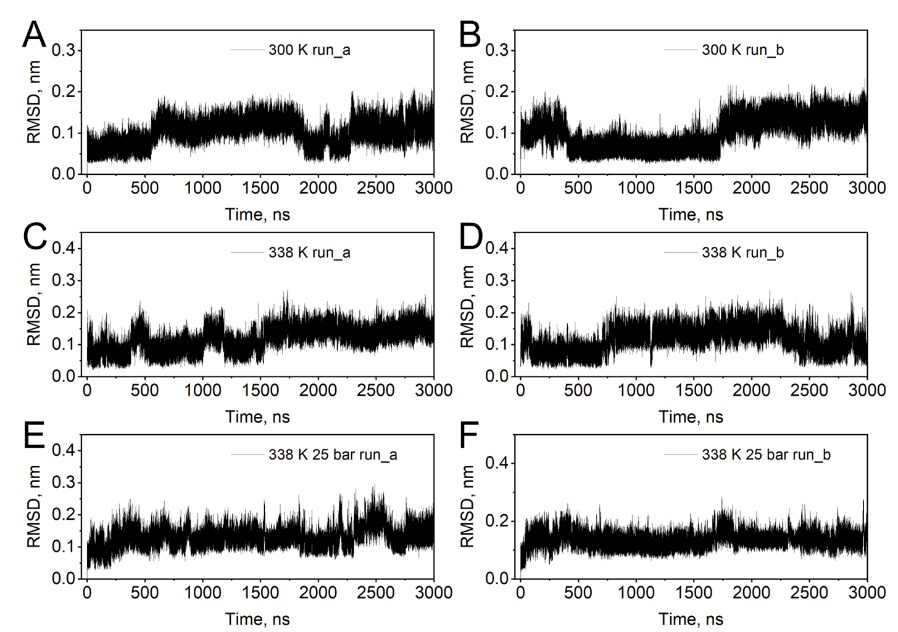

Fig. S8. RMSD of protein C_α_ for residues corresponding to H3 (see *Figure 5 A* in the Main Text) for two independent runs at (a, b) 300 K, (c, d) 338 K, and (e, f) 338 K 25 bar for 3 microseconds.

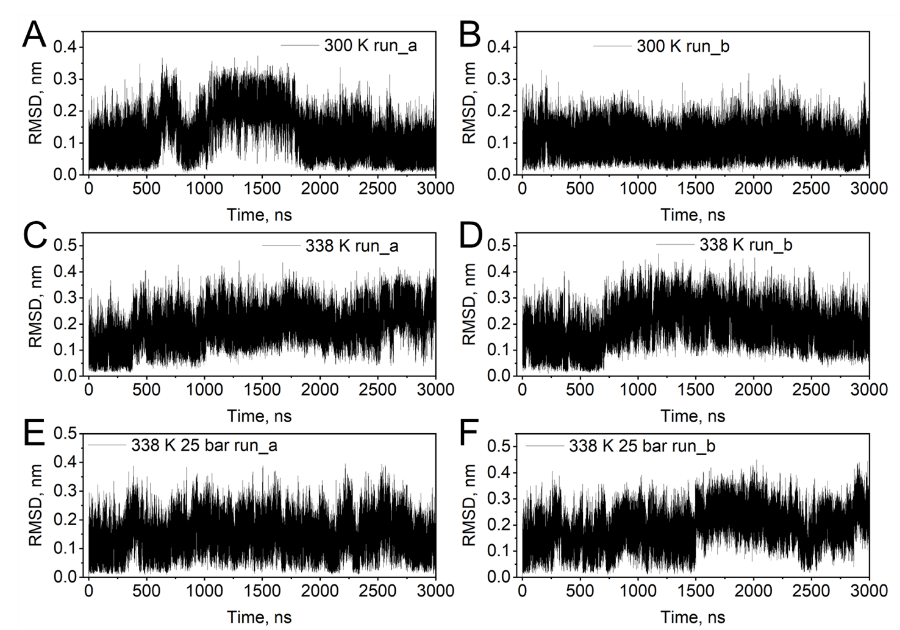

Fig. S9. RMSD of protein C_α_ for residues corresponding to S1 (see *Figure 5 A* in the Main Text) for two independent runs at (a, b) 300 K, (c, d) 338 K, and (e, f) 338 K 25 bar for 3 microseconds.

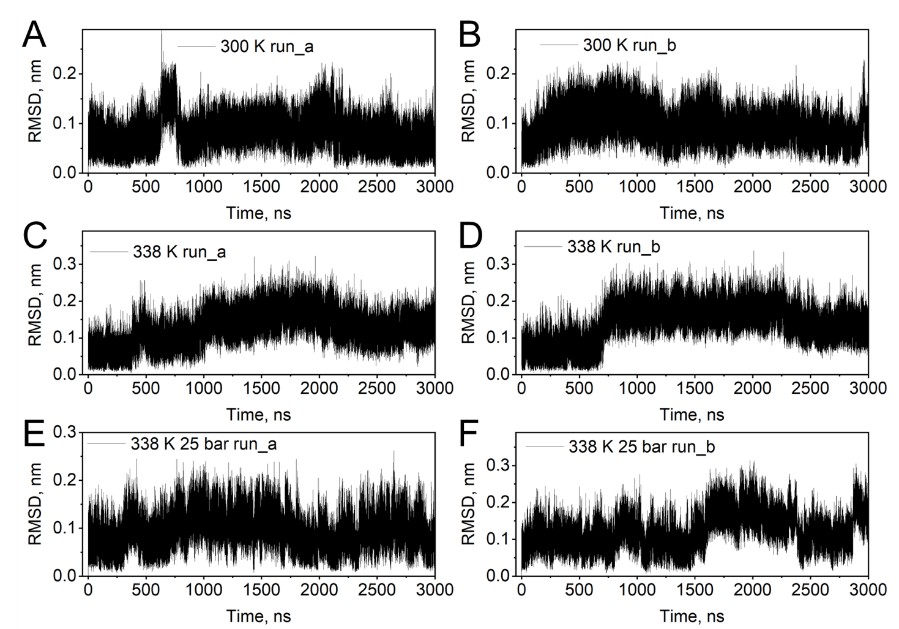

Fig. S10. RMSD of protein C_α_ for residues corresponding to S2 (see *Figure 5 A* in the Main Text) for two independent runs at (a, b) 300 K, (c, d) 338 K, and (e, f) 338 K 25 bar for 3 microseconds.

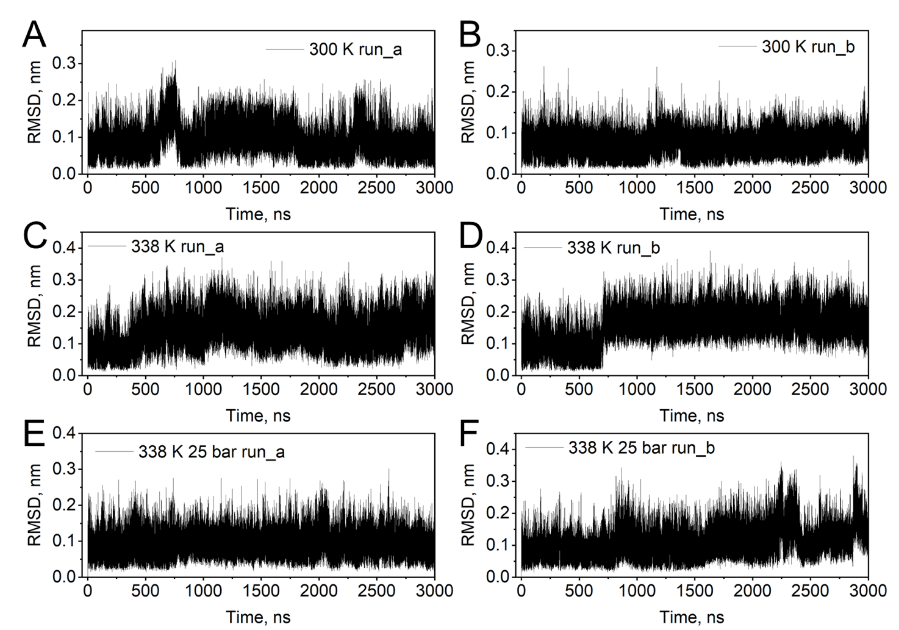

Fig. S11. RMSD of protein C_α_ for residues corresponding to H4 (see *Figure 5 A* in the Main Text) for two independent runs at (a, b) 300 K, (c, d) 338 K, and (e, f) 338 K 25 bar for 3 microseconds.

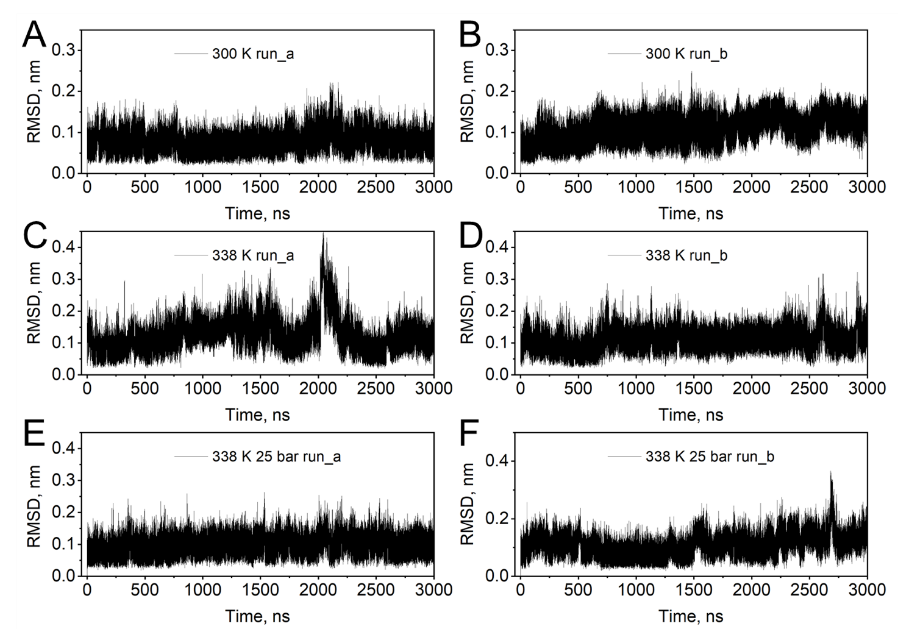

Fig. S12. RMSD of protein C_α_ for residues corresponding to H5 (see *Figure 5 A* in the Main Text) for two independent runs at (a, b) 300 K, (c, d) 338 K, and (e, f) 338 K 25 bar for 3 microseconds.

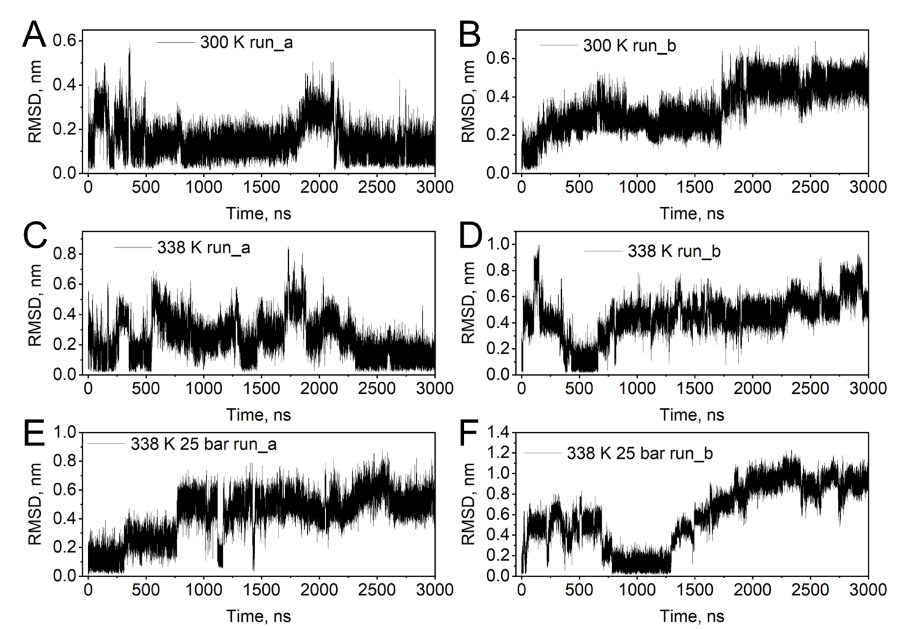

Fig. S13. RMSD of protein C_α_ for residues corresponding to H6 (see *Figure 5 A* in the Main Text) for two independent runs at (a, b) 300 K, (c, d) 338 K, and (e, f) 338 K 25 bar for 3 microseconds.

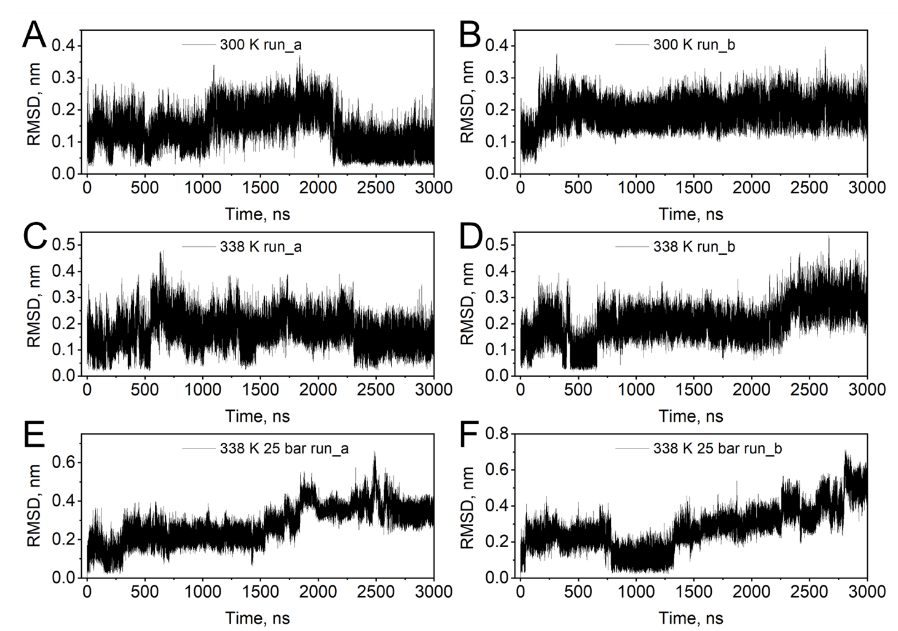

Fig. S14. RMSD of protein C_α_ for residues corresponding to H7 (see *Figure 5 A* in the Main Text) for two independent runs at (a, b) 300 K, (c, d) 338 K, and (e, f) 338 K 25 bar for 3 microseconds.

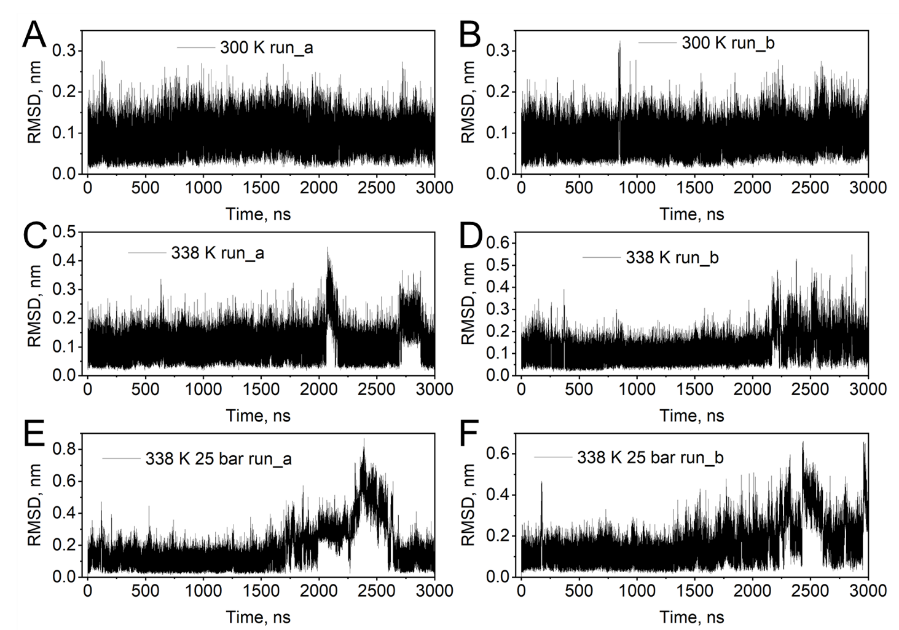

Fig. S15. RMSD of protein C_α_ for residues corresponding to H8 (see *Figure 5 A* in the Main Text) for two independent runs at (a, b) 300 K, (c, d) 338 K, and (e, f) 338 K 25 bar for 3 microseconds.

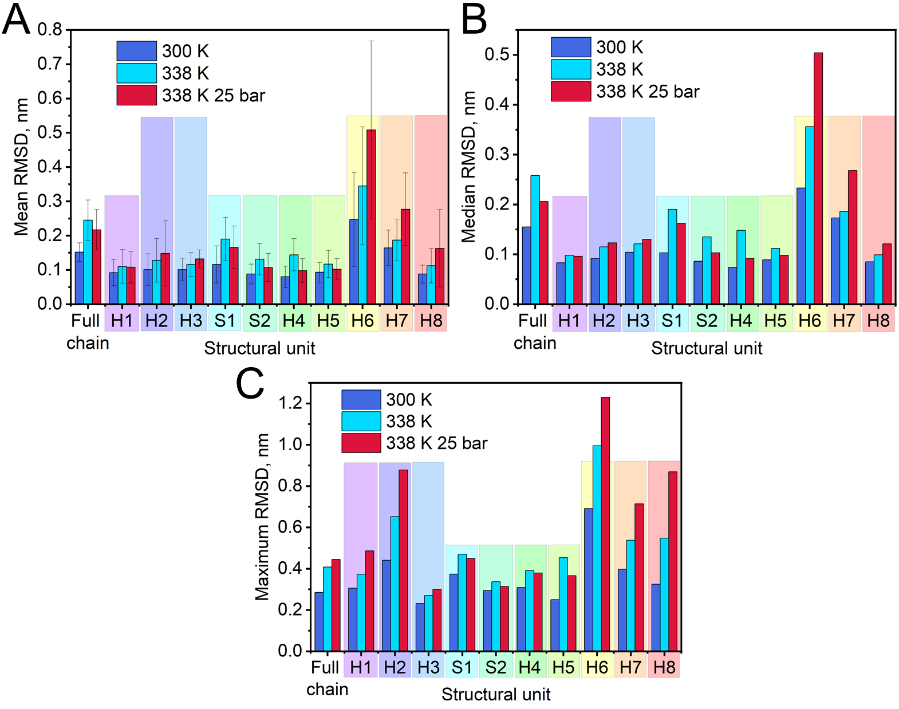

Fig. S16. Summary of RMSD for the lysozyme molecule and its secondary structure units under different conditions (refer to *Table 1* in the Main Text). Rainbow bars show the corresponding color for the unit (referred to in *Figure 5 A* in the Main Text), and their relative height visually represents the impact of pressure on the unit organization compared to the effect of high temperature only (tall bars – pressure disorganizes the unit, short bars – organizes).

Table S1. Relative change (%) in mean, median, and maximum RMSD value (%) for C_α_ by comparing two different states (338 K over 300 K and 338 K 25 bar over 300 K, referred to in Table 1) over two runs for 3 µs (with a step of 1 ps) for whole protein chains and each structural element of its secondary structure (referred to in *Figure 5* *A* in the Main Text). A graphical representation of the RMSD summary is presented in *Fig. S17*.

| **Structure/unit** | **% of change for** | | | | | |
| --- | --- | --- | --- | --- | --- | --- |
|  | **mean RMSD** | | **median RMSD** | | **max RMSD** | |
|  | **338 K**  **over**  **300 K** | **338 K**  **25 bar**  **over**  **300 K** | **338 K**  **over**  **300 K** | **338 K**  **25 bar**  **over**  **300 K** | **338 K**  **over**  **300 K** | **338 K**  **25 bar**  **over**  **300 K** |
| **Entire protein chain** | 61.2 | 42.8 | 66.5 | 32.9 | 43.2 | 56.1 |
| **H1**  RCELAAAMKR | 19.6 | 17.4 | 18.1 | 15.7 | 21.9 | 58.8 |
| **H2**  YRG | 26.7 | 46.5 | 25.0 | 33.7 | 47.8 | 99.1 |
| **H3**  LGNWVCAAKFES | 14.9 | 30.7 | 16.3 | 25.0 | 16.4 | 29.3 |
| **S1**  TNR | 63.8 | 43.1 | 84.5 | 57.3 | 25.7 | 20.3 |
| **S2**  TDY | 48.9 | 21.6 | 57.0 | 19.8 | 14.6 | 6.8 |
| **H4**  CSALL | 80.0 | 22.5 | 100.0 | 24.3 | 26.5 | 22.7 |
| **H5**  TASVNCAKKIV | 25.8 | 9.7 | 25.8 | 10.1 | 81.2 | 46.4 |
| **H6**  GMNA | 39.7 | 105.7 | 52.8 | 116.3 | 44.0 | 78.0 |
| **H7**  VAWRNR | 14.0 | 68.9 | 7.5 | 54.9 | 35.2 | 79.4 |
| **H8**  VQAWI | 27.3 | 85.2 | 16.5 | 42.4 | 68.6 | 167.4 |

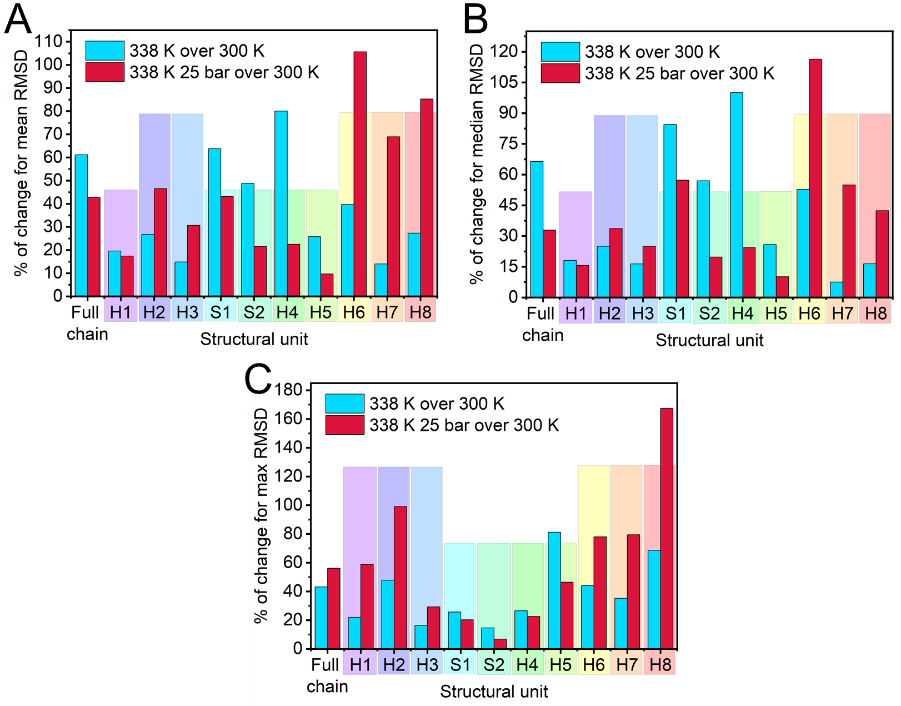

Fig. S17. Summary of the relative change (%) in mean, median, and maximum RMSD value (nm) for C_α_ by comparing two different states (338 K over 300 K and 338 K 25 bar over 300 K, referred to in *Table 1* in the Main Text, *Fig. S16* and Table S1). Rainbow bars show the corresponding color for the unit (referred to in *Figure 5 A* in the Main Text), and their relative height visually shows the impact of pressure on the unit organization, compared to the effect of high temperature only (tall bars – pressure disorganizes the unit, short bars – organizes).

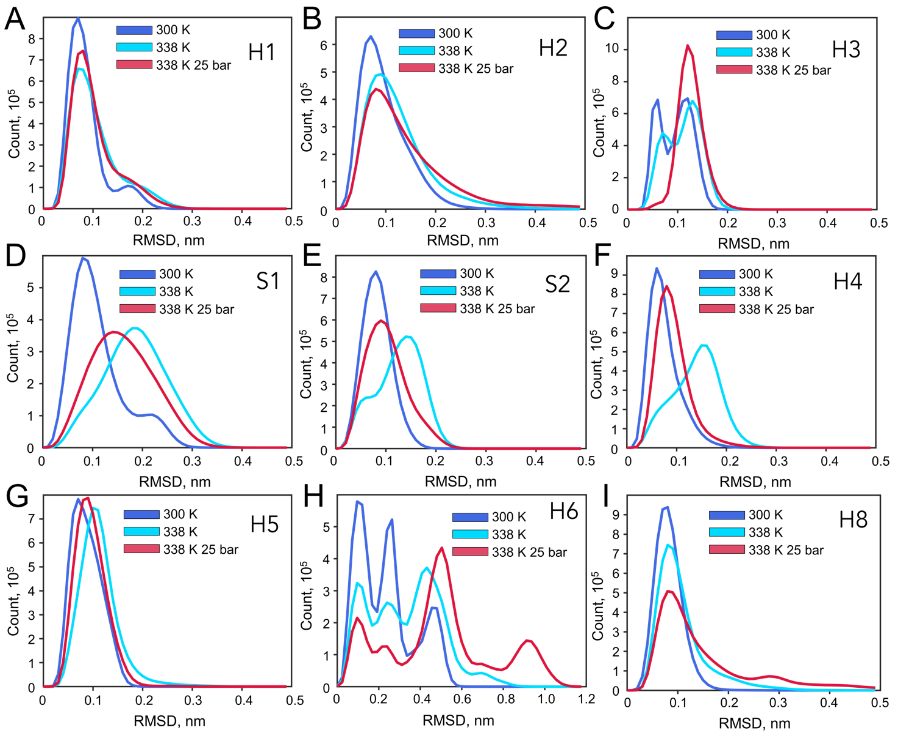

Fig. S18. Diagrams of cumulative (0 – 3000000 ps) RMSD of C_α_ over two runs (referred to in *Figures S4-S14*) for separate units: (a) H1, (b) H2, (c) H3, (d) S1, (e) S2, (f) H4, (g) H5, (h) H6, and (i) H8 under different simulation conditions.

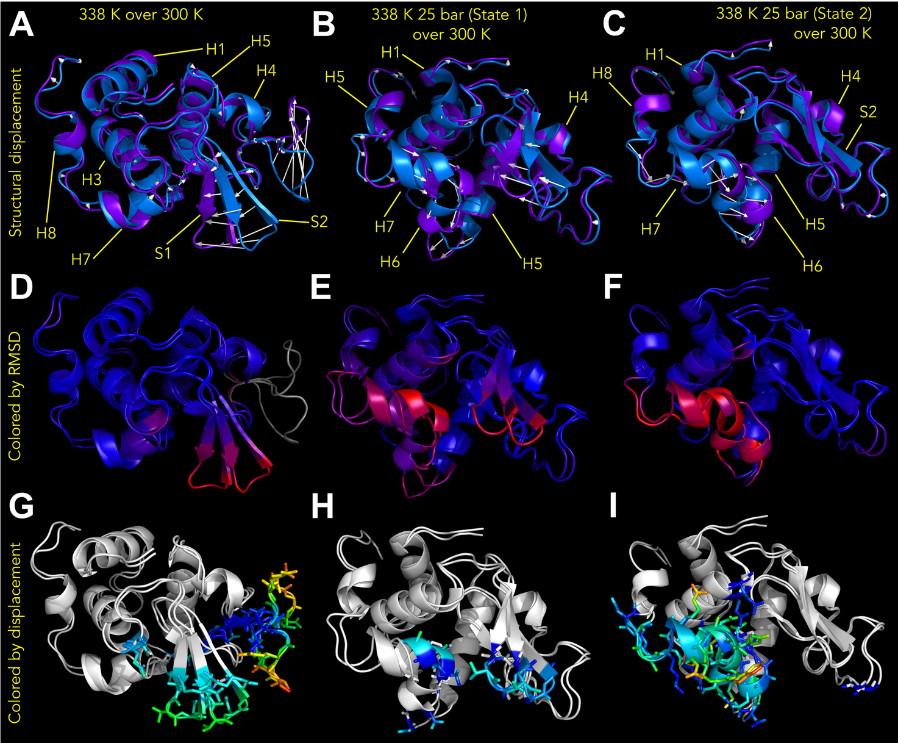

Fig. S19. Pairwise structural alignment between the protein structure at 300 K and the structures at (a, d, g) 338 K, (b, e, h) at 338 K 25 bar (State 1), (c, f, i) at 338 K 25 bar (State 2) and different representation modes: (a–c) coloring by whole structure displacement; arrows represent the displacement vectors of the structure (purple) under different simulation conditions from the initial structure at 300 K (blue); (d–f) coloring by the RMSD value (blue– low RMSD, red– high RMSD, gray– no alignment); (g–i) images show regions in the backbone/sidechain structure that underwent a more significant displacement (in rainbow colors, blue corresponds to a minimum displacement, red – to a maximum) than the other residues in the chain (denoted as white). Visualization: Pymol 1.7/2.5.5.

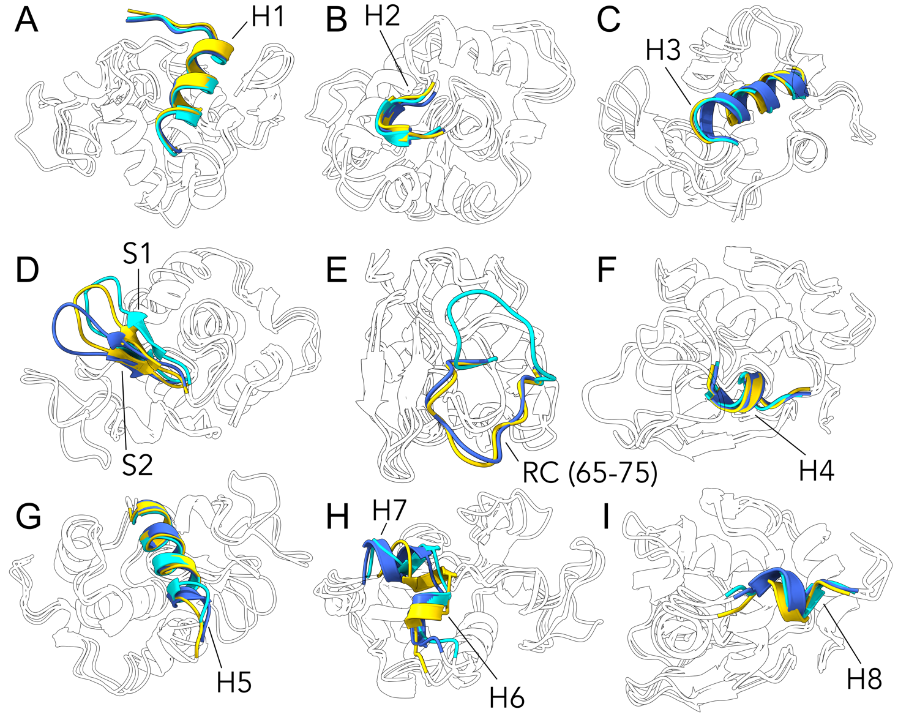

Fig. S20. Pairwise structural alignment between the protein structure at 300 K (blue) with the structures at 338 K (cyan) and at 338 K 25 bar (State 1, yellow). Structural changes of each separate element of the secondary structure of the protein (referred to in *Figure 5 A* in the Main Text) are displayed in corresponding colors (a – H1, b – H2, c – H3, d – S1 and S2, e – a segment of the random coil sequence, displaying maximum displacement, f – H4, g – H5, h – H6, and H7, i – H8), whereas the other chain is shown as silhouettes. Visualization: ChimeraX 1.7.

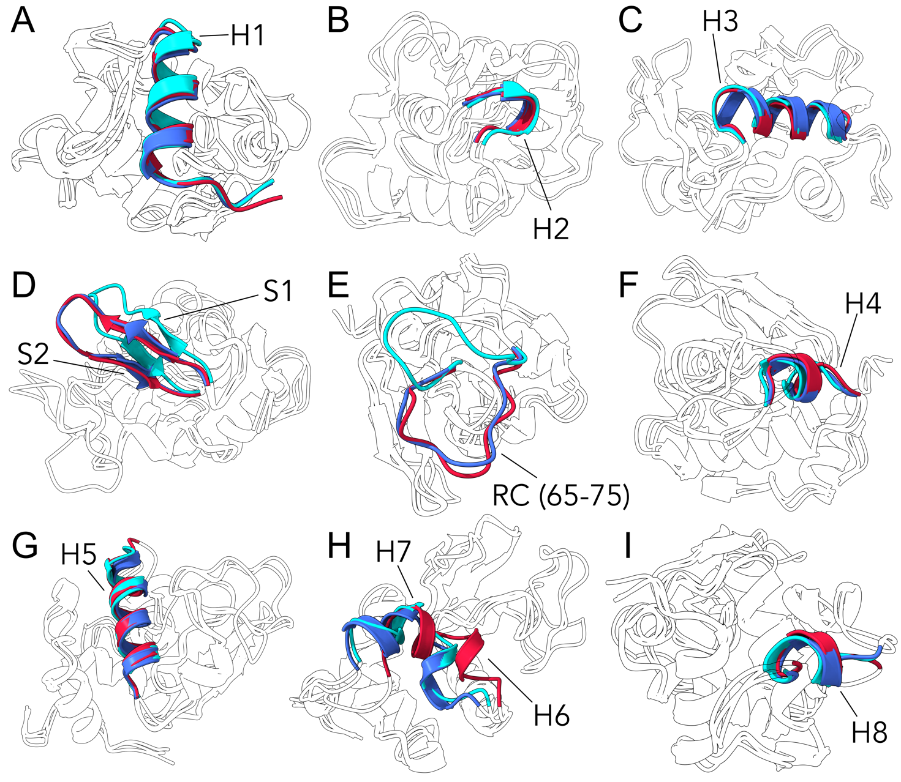

Fig. S21. Pairwise structural alignment between the protein structure at 300 K (blue) with the structures at 338 K (cyan) and at 338 K 25 bar (State 2, red). Structural changes of each separate element of the secondary structure of the protein (referred to in *Figure 5 A* in the Main Text) is displayed in the corresponding color (a – H1, b – H2, c – H3, d – S1 and S2, e – a segment of the random coil sequence, displaying the maximum displacement, f – H4, g – H5, h – H6 and H7, i – H8), whereas the other chain is shown as silhouettes. Visualization: ChimeraX 1.7.

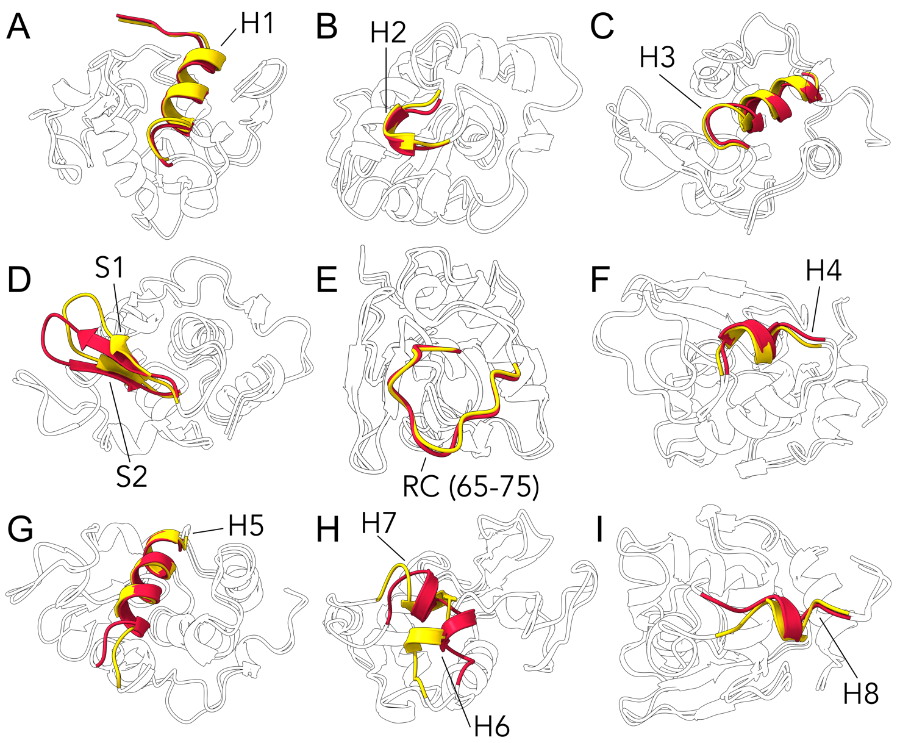

Fig. S22. Pairwise structural alignment between the protein structure at 338 K 25 bar (State 1, red) and the structure at 338 K 25 bar (State 2, yellow) represents two temporally separated states at 327309 and 1843799 ps. Structural changes of each separate element of the secondary structure of the protein (referred to in *Figure 5 A* in the Main Text) are displayed in the corresponding color (a – H1, b – H2, c – H3, d – S1 and S2, e – a segment of the random coil sequence, displaying maximum displacement, f – H4, g – H5, h – H6 and H7, i – H8), whereas the other chain is shown as silhouettes. Visualization: ChimeraX 1.7.

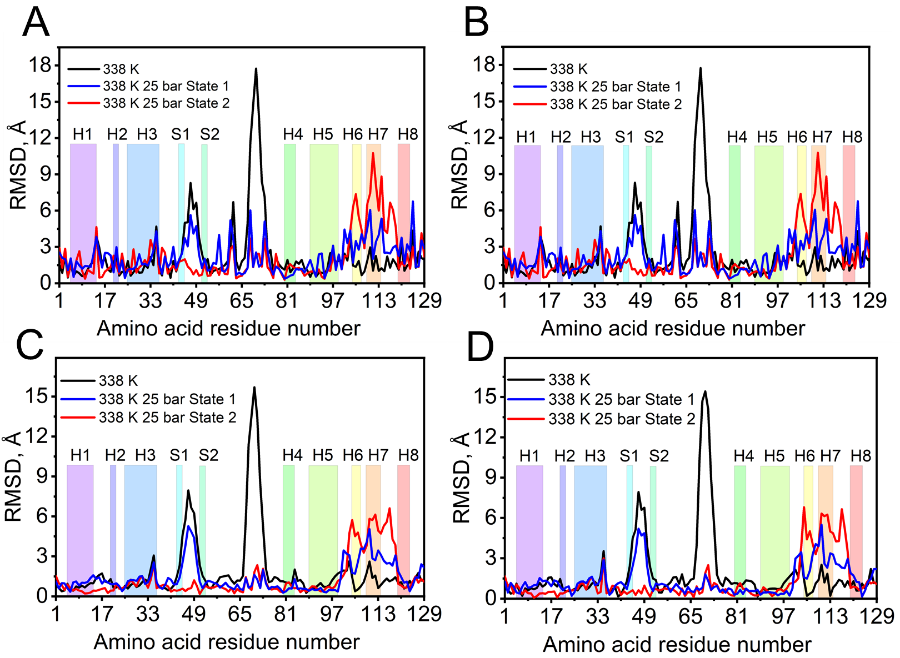

Fig. S23. Pairwise RMSD per residue between the single-point structures at 338 K and 338 K 25 bar (two temporally separated states at 327309 and 1843799 ps) and a structure at 300 K: (a) RMSD by all atoms, (b) minimum RMSD value by all atoms, (c) RMSD by C_α_, and (d) RMSD by backbone structure (referred to in *Fig. S19*).

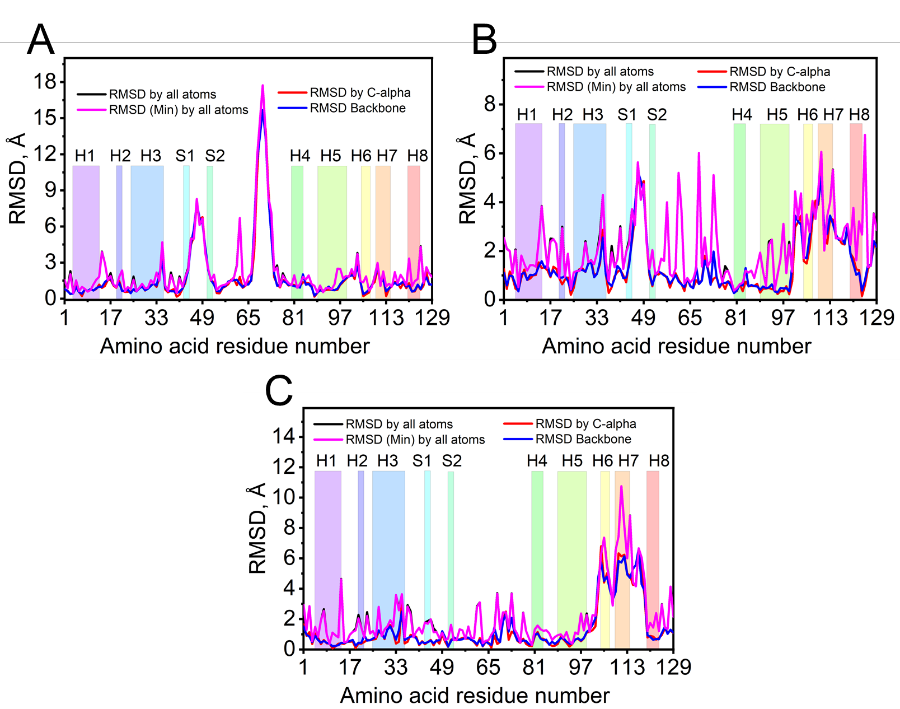

Fig. S24. Another kind of representation of the data is shown in *Fig. S23*: pairwise RMSD per residue calculated by all atoms, by C_α_ and by the backbone between the single-point structures (a) at 338 K, (b) at 338 K 25 bar (state 1), and (c) at 338 K 25 bar (state 2) regarding the structure at 300 K (referred to in *Fig. S19*).

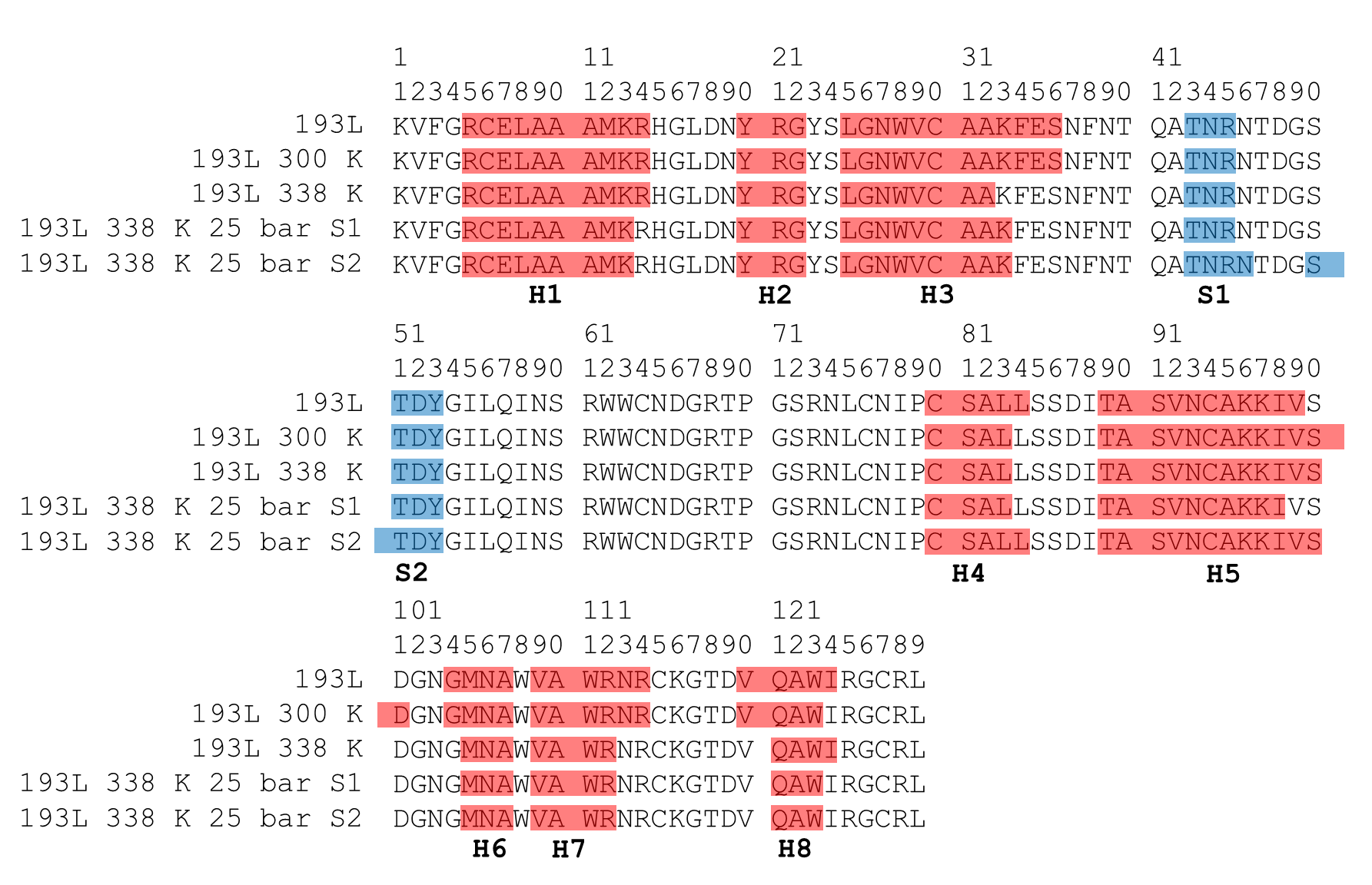

Fig. S25. Secondary structure similarity of 5 protein models under different simulation conditions: the original one (PDB ID: 193l), at 300 K, at 338 K, at 338 K 25 bar (state 1), and at 338 K 25 bar (state 2). Red regions display positions of specific amino acids that adopt the α-helix conformation, whereas the blue regions correspond to β-sheet structures. Secondary structure analysis was performed using Chimera 1.18 (according to DSSP algorithm).

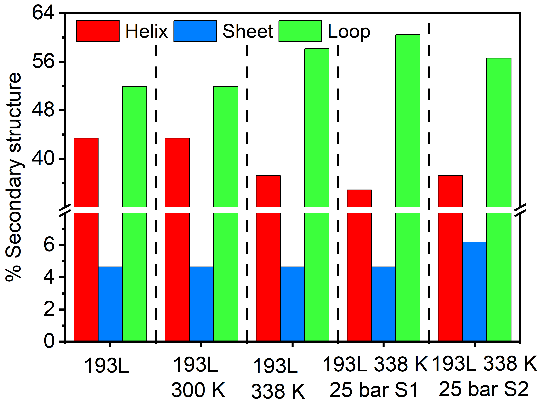

Fig. S26. Graphical representation of the secondary structure composition of the HEWL protein under different simulation conditions (referred to in *Fig. S25* and *Table S2*).

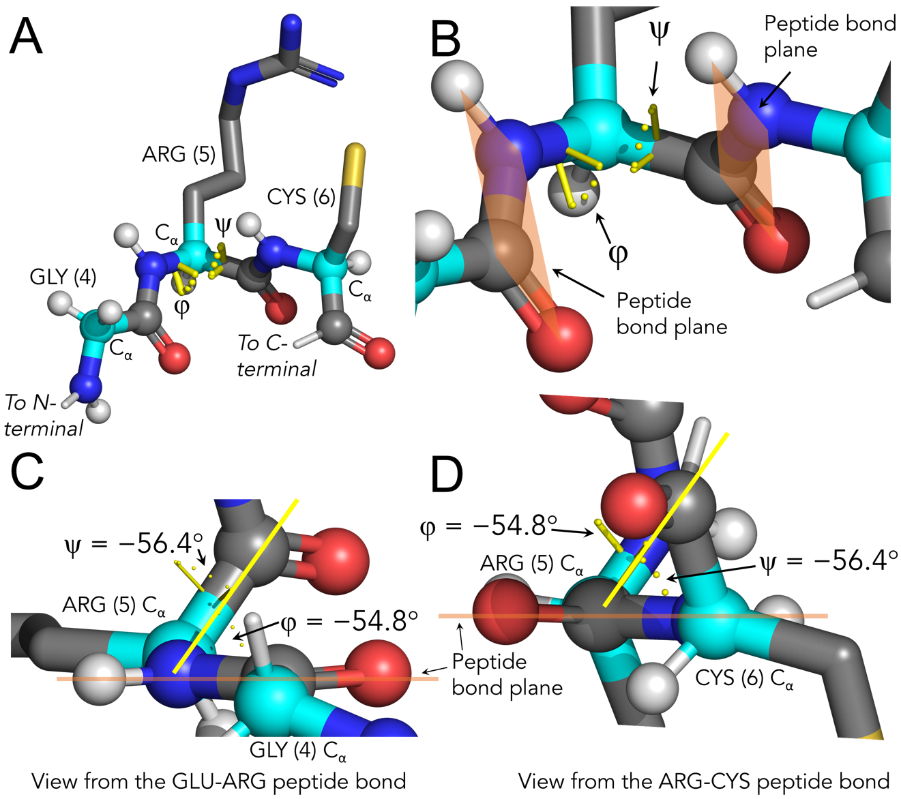

Fig. S27. Illustrative figures for the φ and ψ angle positions in the protein (PDB:193L) fragment (4-6: GLY-ARG-CYS): (a) general and (b) a detailed view of the sequence fragment indicating the key elements of the structure required to calculate the corresponding angle values and (c, d) different views of the fragment showing the exact positions of the angles. Visualization: Pymol 1.7/2.5.5.

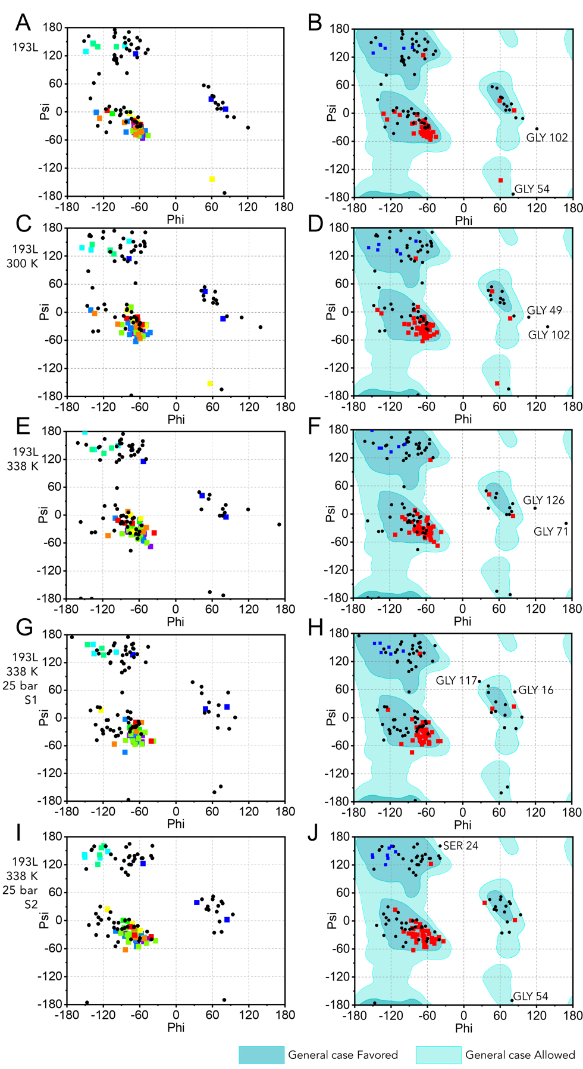

Fig. S28. Ramachandran plots of HEWL under different simulation conditions: (a, b) original structure (PDB ID: 193l), (c, d) at 300 K, (e, f) at 338 K, (g, h) at 338 K 25 bar (state 1), and (i, j) at 338 K 25 bar (state 2). Left panel (a, c, e, g, i) shows the distribution of the dihedral angles of the amino acid residues: colored squares denote the structural elements depicted in *Figure 5 A* (black rounds denote loops) considering also the secondary structure changes presented in *Fig. S25*. Right panel (b, d, f, h, j) the same data points but showing the distribution of α-helixes (red squares), β-sheets (blue squares), and loops (black rounds) on the background of a general case of the distribution of dihedral angles. Points inside the favored regions denote relaxed angles, whereas points inside of the allowed region and outside of it denote strained angles.

Table S2. Secondary structure composition of the HEWL under different simulation conditions (referred to in *Fig. S25*).

| **Structure** | **Number of residues in unit** | | | **Secondary structure, %** | | |
| --- | --- | --- | --- | --- | --- | --- |
|  | **Helix** | **Sheet** | **Loop** | **Helix** | **Sheet** | **Loop** |
| **193L** | 56 | 6 | 67 | 43.41 | 4.65 | 51.94 |
| **193L 300 K** | 56 | 6 | 67 | 43.41 | 4.65 | 51.94 |
| **193L 338 K** | 48 | 6 | 75 | 37.21 | 4.65 | 58.14 |
| **193L 338 K 25 bar S1** | 45 | 6 | 78 | 34.88 | 4.65 | 60.47 |
| **193L 338 K 25 bar S2** | 48 | 8 | 73 | 37.21 | 6.20 | 56.59 |

**SI References**

1. F. Oosawa, M. Kasai, A theory of linear and helical aggregations of macromolecules. Journal of Molecular Biology 4, 10–21 (1962).
2. S. I. A. Cohen, et al., Nucleated polymerization with secondary pathways. I. Time evolution of the principal moments. J. Chem. Phys. 135, 065105 (2011).
3. S. I. A. Cohen, et al., Nucleated polymerization with secondary pathways. II. Determination of self-consistent solutions to growth processes described by non-linear master equations. J. Chem. Phys. 135, 065106 (2011).

4. S. I. A. Cohen, et al., Nucleated polymerization with secondary pathways. III. Equilibrium behavior and oligomer populations. J. Chem. Phys. 135, 065107 (2011).
5. T. C. T. Michaels, et al., Amyloid formation as a protein phase transition. Nat Rev Phys 5, 379–397 (2023).
6. A. J. Dear, et al., Kinetic diversity of amyloid oligomers. Proc. Natl. Acad. Sci. U. S. A. 117, 12087–12094 (2020).
7. S. Roy, N. V. Bapat, J. Derr, S. Rajamani, S. Sengupta, Emergence of ribozyme and tRNA-like structures from mineral-rich muddy pools on prebiotic earth. J. Theor. Biol. 506, 110446 (2020).
8. M. Jafari, F. Mehrnejad, Molecular insight into human lysozyme and its ability to form amyloid fibrils in high concentrations of sodium dodecyl sulfate: A view from molecular dynamics simulations. PLoS One 11, e0165213 (2016).
